# Biologic that disrupts PDE11A4 homodimerization in hippocampus CA1 reverses age-related proteinopathies in PDE11A4 and cognitive decline of social memories

**DOI:** 10.1101/2022.08.31.506102

**Authors:** Katy Pilarzyk, Will Capell, Audrey Rips-Goodwin, Latarsha Porcher, Michy P. Kelly

## Abstract

Age-related proteinopathies in phosphodiesterase 11A (PDE11A), an enzyme that degrades 3’,5’-cAMP/cGMP and is enriched in the ventral hippocampal formation (VHIPP), drive age-related cognitive decline (ARCD) of social memories. In the VHIPP, age-related increases in PDE11A4 occur specifically within the membrane compartment and ectopically accumulate in filamentous structures termed ghost axons. Previous *in vitro* studies show that disrupting PDE11 homodimerization by expressing an isolated PDE11A-GAFB domain, which acts as a “negative sink” for monomers, selectively degrades membrane-associated PDE11A4 and prevents the punctate accumulation of PDE11A4. Therefore, we determined if disrupting PDE11A4 homodimerization *in vivo* via the expression of an isolated PDE11A4-GAFB domain would be sufficient to reverse 1) age-related accumulations of PDE11A4 in VHIPP ghost axons and 2) ARCD of social memories. Indeed, *in vivo* lentiviral expression of the isolated PDE11A4-GAFB domain in hippocampal CA1 reversed the age-related accumulation of PDE11A4 in ghost axons, reversed ACRD of social transmission of food preference memory (STFP), and improved remote long-term memory for social odor recognition (SOR) without affecting memory for non-social odor recognition. *In vitro* studies suggest that disrupting homodimerization of PDE11A4 does not directly alter the catalytic activity of the enzyme but may reverse age-related decreases in cGMP by dispersing the accumulation of the enzyme independently of other intramolecular mechanisms previously established to disperse PDE11A4 (e.g., phosphorylation of PDE11A4 at serine 162). Altogether, these data suggest that a biologic designed to disrupt PDE11A4 homodimerization may serve to ameliorate age-related deficits in hippocampal cyclic nucleotide signaling and subsequent ARCD of remote social memory.

## INTRODUCTION

Decreases in 3’,5’-cyclic nucleotide signaling in the aged and demented hippocampus contribute to cognitive decline and are related, in part, to changes in the enzymes that break down cyclic nucleotides [4-6]. There are eleven families of 3’,5’-cyclic nucleotide phosphodiesterases (PDEs) that are the only known enzymes to hydrolyze 3’,5’-cyclic adenosine monophosphate (cAMP) and 3’,5’-cyclic guanosine monophosphate (cGMP), and their expression across subcellular compartments differs between families [7]. For example, some PDEs are expressed more in the cytosol than the membrane (e.g., PDE11A), while others are more highly expressed in the membrane versus cytosol (including PDE2A, PDE9A and PDE10A [8, 9]). Due to the fact that PDEs are localized to specific subcellular domains, they are able to regulate individual pools or nanodomains of cyclic nucleotide signaling [7]. Such subcellular compartmentalization of cyclic nucleotide signaling allows a single cell to respond specifically to simultaneous intra- and/or extracellular signals. Therefore, the subcellular localization of any PDE is equally important to its actual catalytic activity when considering its function. PDEs can become overexpressed and/or mislocalized with age and/or disease, which compromises the integrity of this physiological segregation of signals [9-12]. Indeed, age-related diseases and neuropsychiatric diseases can show a loss of cyclic nucleotide signaling in one subcellular compartment but not another [5, 6, 13-16], suggesting therapeutic strategies should optimally target enzymes in a compartment-specific manner.

Of the eleven families of PDEs, phosphodiesterase 11A (PDE11A) has garnered particular interest in the context of altered cyclic nucleotide signaling related to ARCD and Alzheimer’s disease [2, 12, 17, 18]. PDE11A is encoded by a single gene and has four isoforms [9-12, 14]. While protein for PDE11A4—the isoform expressed in brain—is found across all subcellular compartments, it is particularly enriched in the cytosolic versus membrane and nuclear compartments [8]. The PDE11A catalytic domain is located within the C-terminal region, which is common to all isoforms, while the N-terminal region serves a regulatory function and is unique to each isoform (c.f., [8, 19]). The regulatory N-terminus of PDE11A4, the longest PDE11A isoform, is unique in that it contains two full GAF (cGMP binding PDE, Anabaena adenylyl cyclase and *E. coli* FhlA) domains. The GAF-A domain binds cGMP as a potential allosteric regulatory site and the GAF-B domain regulates protein-protein interactions, including homodimerization [1, 20, 21]. PDE11A4 is unique in that it is the only PDE whose expression in brain emanates solely from the extended hippocampal formation, a brain region critical to learning and memory [12, 22] and vulnerable to age-related deficits in cyclic nucleotide signaling [6]. Possibly contributing to these hippocampal cyclic nucleotide signaling deficits are age-related increases in PDE11A4 expression that are conserved across mice, rats and humans [2, 12]. These age-related increases in PDE11A4 protein expression are deleterious as 1) *Pde11a* KO mice are protected against age-related cognitive decline (ARCD) of remote social associative long-term memories (aLTMs) [2] and 2) mimicking age-related overexpression of PDE11A4 in the CA1 field of hippocampus of either young or old *Pde11a* KO mice is sufficient to trigger decline of remote social aLTMs [2, 23].

While PDE11A4 protein expression in the young adult hippocampus is significantly higher in the cytosolic versus membrane compartment, age-related increases in PDE11A4 are found specifically in the membrane compartment of the ventral hippocampal formation (VHIPP; [2]). Further, aging triggers a punctate accumulation of PDE11A4 protein in filamentous structures termed ghost axons (i.e., in homage to tau ghost tangles; [2]) that are rarely seen in the young brain. Therefore, these age-related increases in membrane-associated PDE11A4 and PDE11A4 ghost axons may be considered ectopic. That said, the membrane pool of PDE11A4—although lesser in relative quantity—continually shows itself to be critical in regulating social behaviors [3, 18, 24]. For instance, we found that social isolation selectively decreases expression of membrane-associated PDE11A4 in the VHIPP, and that these isolation-induced decreases in PDE11A4 are sufficient to cause changes in subsequent social preferences and social memory [18, 24]. Similarly, PDE11A4 protein expression differences in VHIPP of BALB/cJ versus C57BL/6J mice are restricted to the membrane pool [1]. Of direct relevance to the current study, the increased protein expression of membrane-associated PDE11A4 found in the VHIPP of BALB/cJ vs C57BL/6J mice appears to be driven by a single point mutation at amino acid 499 within the GAF-B domain, which strengthens the homodimerization and punctate accumulation of PDE11A4 [1, 3]. Interestingly, disrupting PDE11A4 homodimerization *in vitro* by expressing an isolated GAF-B domain that acts as a negative sink disperses the punctate accumulation of PDE11A4 and selectively reduces expression of membrane-associated PDE11A4 [1]. This suggests that disrupting PDE11A4 homodimerization *in vivo* may represent a therapeutic option capable of treating age-related increases in PDE11A4 expression in a compartment-specific manner and, thus, ARCD of social memories. Indeed, we previously showed that social preference of C57BL/6J mice can be altered by manipulating PDE11A4 homodimerization selectively within the CA1 field of hippocampus [3]. Therefore, we seek to determine if a biologic that decreases PDE11A4 homodimerization will be sufficient to reverse age-related accumulations of PDE11A4 in ghost axons and rescue ARCD of social memories.

## METHODS

### Subjects

C57BL6/J mice were originally obtained from Jackson Laboratory (Bar Harbor, ME) and the line was maintained at the University of South Carolina. As previously described [3, 23, 25], the *Pde11a* mouse line obtained from Deltagen (San Mateo, CA) was maintained on a mixed C57BL6 background (99.8% multiple C57BL/6 substrains, 0.2% 129P2/OlaHsd) [3]. *Pde11a* mice were bred at the University of South Carolina in heterozygous (HT) x HT trio-matings. Multiple same-sex wild-type (WT) mice along with one HT mouse (used as the demonstrator in social transmission of food preference—see more below) were weaned and caged together. Typically, cages included ∼4-5 mice/cage. We do not believe litter effects are driving findings here due to each cohort being composed of WT mice from multiple litters and, in the case of NSOR and SOR, the datasets are composed of multiple cohorts born and tested at different times [3, 26]. While both males and females were used in experiments, we are underpowered to analyze for sex effects (see figure legends for specific *n*’s/sex/group/experiment). In these studies, young mice were defined as 2-6 months and old mice were defined as 18-24 months. “Young” mice included both young *Pde11a* WT mice surgerized alongside old *Pde11a* WT mice (i.e., receiving bilateral injections of mCherry lentivirus to the dorsal and ventral hippocampi) and unsurgerized young C57BL6/J mice that were used as an internal control for the assays (see figures 2A, 2B). Since no obvious differences were found between groups of young surgerized *Pde11a* WT mice and young unsurgerized C57BL6/J mice, the data from these 2 subgroups were subsequently combined into a singular “young” group (see figures 2C-G). All mice used in experiments were generally healthy throughout the duration of testing. As previously described [26], we do not conduct gross pathology but mice are routinely assessed by husbandry, veterinary, and laboratory staff. Mice with lethargy, altered gait, signs of malnutrition or dehydration, tumors >1 cm, are removed from study and euthanized. We aim to study the effects of healthy aging [26], therefore if upon brain dissection we had found evidence of an anatomical abnormality (e.g., a pituitary tumor), the animals would have been excluded from the study (note: no occurrences in this study). A 12:12 light:dark cycle and ad lib access to food and water were provided. Experiments were conducted in accordance with the National Institutes of Health Guide for the Care and Use of Laboratory Animals (Pub 85-23, revised 1996) and were fully approved by the Institutional Animal Care and Use Committee of the University of South Carolina.

### Plasmid generation

Plasmids were generated as previously described [1, 2]. Briefly, constructs were generated by Genscript (Piscataway, NJ) that expressed either mCherry alone, an mCherry-tagged PDE11A4-GAF-B domain (aa388-558 of NP_001074502.1,which includes 14 upstream amino acids as a spacer), EmGFP alone containing an A206Y mutation to prevent EmGFP dimerization [27], or an EmGFP-tagged mouse *Pde11a4* (NM_001081033). These constructs were initially generated on a pUC57 backbone and then subcloned into a pcDNA3.1+ mammalian expression vector (Life Technologies; Walthan, MA). The QuickChange procedure/products were used to generate *Pde11a4* mutations as per manufacturer’s instructions (Agilent Technologies; Santa Clara, CA). Oligonucleotide primers were generated by Integrated DNA Technologies (Coralville, IA) and mutated DNA sequences were verified by Functional Biosciences (Madison, WI).

### Cell culture and transfections

COS-1 (male line), HEK293T (female line), and HT-22 (sex undefined) cell culture and transfection were performed as previously described [1]. While kept in t-75 flasks, cells were grown in Dulbecco’s Modified Eagle Medium (DMEM)with (HT-22) or without (COS-1, HEK293T) sodium pyruvate (GIBCO; Gaithersburg, MD; or Corning, Manassas, VA,), 10% fetal bovine serum (FBS), and 1% Penicillin/Streptomycin (P/S) (GE Healthcare Life Sciences; Logan, UT, USA). Cells were incubated at 37°C/5% CO_2_ and passaged using TrypLE Express (GIBCO; Gaithersburg, MD, USA) as a dissociation agent once 70-90% confluent. One day before transfection began, cells were plated along with DMEM+FBS+P/S in either 24-well plates for imaging or 100 mm dishes for biochemistry. The day of transfection, Optimem (GIBCO) replaced the media. According to manufacturer protocol (ratio of 3.75 ug DNA plus 10uL lipofectamine per 10 mLs of media), cells were transfected with Lipofectamine 2000 (Invitrogen; Carlsbad, CA). ∼19 hours post-transfection (PT), the Optimem/Lipofectamine solution was removed and replaced with DMEM+FBS+P/S. Normally, cells grew for five hours before sample processing, which meant they were harvested about 24 hours following transfection. Over the course of experiments, cells were sporadically tested for mycoplasma, with negative results always obtained. For assessment of subcellular trafficking, paraformaldehyde (4%) in PBS was used to fix the cells for fifteen minutes, after which they were kept in PBS until imaging. Images were captured using NIS-Elements BR-2.3 (Nikon,Tokyo Japan) on a CoolSNAP EZ CCD camera (Photometrics, Tuscon AZ) mounted on an inverted Leica (Wetzlar DE) DMIL microscope with a Fluotar 10X/0.3 ∞/ 1.2 objective.

All images pertaining to an experiment were quantified by an experimenter blind to treatment using the same computer within the same position in the room, the same lighting conditions, and the same percent zoom. Images were loaded onto a gridded template to facilitate keeping track of count locations within the image, and an experimenter scored each image box by box, with cells along the top and left edges of the image as a whole not included to follow stereological best practices. Images were quantified in a counterbalanced manner such that 1 picture from each condition was evaluated before moving onto a 2_nd_ image from that condition. The experimenter classified cells as exhibiting either cytosolic-only labeling or punctate labeling (with or without cytosolic labeling present), with data expressed as the % of the total number of labeled cells that exhibited punctate labeling.

### Stereotaxic Surgeries

Stereotaxic surgeries and injections were performed, as previously described [3, 23], using a NeuroStar motorized stereotaxic, drill, and injection robot (Tubingen, Germany). Mice were anesthetized with a steady flow of oxygen and isoflurane. The mice were induced at 3% isoflurane and maintained at 1-1.5%. Lack of reflexes was verified and the scalp was then shaved and cleaned with betadine. A small incision was made in the scalp and the skull was cleared with sterile saline. Cotton swabs were again used to visualize the skull and locate Bregma. Using the robotic drill, small holes were made above the dorsal and ventral hippocampi as per the following coordinates relative to Bregma: dCA1 AP, −1.7, dCA1 ML, +/-1.6, vCA1 AP, −3.5, vCA1 ML, +/-3.0. At a speed of 10 mm/sec, a Hamilton syringe (custom needle #7804-04: 26s gauge, 1” length, 25 degree bevel) was then then placed to the following depths relative to Bregma: dCA1 DV, 1.3, vCA1 DV, 4.4. After a thirty second pause following the needle movement, the injection robot was used to inject 2 µl of lentivirus at 0.167 µl/minute. Following injection completion, the experimenter waited two minutes to allow the lentivirus to diffuse away from the needle and the needle was raised at the same speed. After all injections were complete, pronged tweezers were used to close the scalp and secured using glutures. Buprenorphine in sterile saline at a dose of 0.1mg/kg was injected IP for pain management. For recovery, the mouse was placed on a warm Deltaphase pad and allowed to recover until moving normally and posturing upright. Mice were returned to their home cages and allowed at least 2 weeks of recovery prior to behavioral testing.

As previously described [3], a lentivirus carrying an mCherry-tagged PDE11A4-GAFB domain served to disrupt PDE11A4 homodimerization, while an mCherry-only virus was used as a negative control. A lentiviral construct was used here in order to compare to previous studies examining the effects of overexpressing PDE11A4 *in vivo* [2, 23], which required the use of a lentiviral cassette due to the large size of PDE11A4. The Viral Vector Core of the University of South Carolina generated the lentiviruses. The viruses were made on an “SPW” backbone that drives expression using the phosphoglycerate kinase 1 (PGK) promoter, which in theory is a ubiquitous promoter and yet is taken up preferentially by neurons [23]. We previously showed that the isolated GAF-B construct disrupts PDE11A4 homodimerization by binding to PDE11A4 monomers and acting as a negative sink that prevents monomers from binding to each other [1]. For reasons that are not well understood, preventing homodimerization via this method triggers proteolytic degradation of membrane-associated PDE11A4 [1]. The lentiviruses were prepared and diluted in 0.2M sucrose/42 mM NaCl/0.84 mM KCl/2.5 mM Na2HPO4/0.46 mM KH2PO4/0.35 mM EDTA and the original titers were as follows: mCherry, 7.37×10E10/mL; GAF-B, 1.82×10E10/ml. Pilot studies using wide-field fluorescent microscopy determined diluting the mCherry-only virus to one-third the original concentration yielded comparable mCherry expression between viruses, and so this concentration was used in experiments. While we found high viral expression throughout CA1 of hippocampus in all mice, a subset of mice exhibited some viral expression in dentate gyrus, CA2, and/or CA3. We do not believe this ectopic viral expression confounds our results given that 1) it is found in both the mCherry-only and mCherry-GAFB groups and 2) PDE11A4 expression does not emanate from the dentate, CA2 nor CA3. Of note, we found that cells in proximal CA1 (closer to CA3) relative to distal CA1 (closer to subiculum) more readily took up the virus as previously noted [23], which mirrors the preferential distribution pattern of endogenous PDE11A4 across CA1 [22]. Importantly, we found no gross cellular toxicity or morphological damage with either virus and animals were healthy following surgery. The accuracy of the injection and expression of the viral construct was verified by direct visualization of raw florescence and/or following amplification of the signal via immunofluorescence (see below).

### Tissue/slide Collection

As previously described (e.g., [23]), mice were euthanized (during light cycle) via rapid cervical dislocation and brains were immediately collected and flash frozen in 2-methylbutane sitting on dry ice. Brain tissue was then stored at −80 °C until cryosectioning at −18 °C. 20 µm sections were collected and thaw mounted onto glass +/+ slides. Slides were briefly dried at room temperature and then temporarily stored in the cryostat until all sections were collected. Once all sections were collected for a given brain, slides were boxed and stored at −80 °C.

### Immunofluorescence

As previously described [22], slides were fixed in room temperature 4% paraformaldehyde in phosphate-buffered saline (PBS) for 20 minutes. After fixation, 3×10 minute washes with PBS and 3×10 minute washes with PBT (phosphate-buffered saline/0.4% BSA/0.3% Triton-X 100) were performed to reduce background. Primary antibodies were combined in a tube with PBT at validated and optimized concentrations: PDE11A4 (Aves custom PDE11A4 #1 at 1:10,000; Fabgennix PPD11A-140AP at 1:1000; Fabgennix PPD11A-150AP at 1:500) and mCherry (ThermoFisher #PA534974 at 1:1000; Invitrogen #M11217 at 1:500; PhosphoSolutions #1203-mCherry at 1:10,000). Multiple PDE11A antibodies were utilized to discern the ectopic accumulation of PDE11A4 in ghost axons. While the PDE11A4#1 antibody labels all PDE11A4 and, thus, detects both diffusely localized and accumulated PDE11A, the PDE11A4-140 and 150 antibodies specifically labels PDE11A4 phosphorylated at serines 117 and/or 124 and, thus, only detects PDE11A4 accumulated in ghost axons [2]. To limit non-specific labeling of the mCherry antibodies hosted in a rodent species (see above Invitrogen and PhosphoSolutions), tissue was pretreated with anti-Mouse FabFragments (0.15mg/ml; Jackson Immunoresearch # 715-007-003) in PBS for 2 hours, followed by 3×10 minute washes in PBT, prior to adding primary antibody. Primary antibody solution was added over brain sections and the slides were kept level at 4°C overnight. Primary antibodies were removed using 4×10 minute washes in PBT. For optimal labeling, PDE11A4 secondary antibody (Alexafluor 488 AffiniPure Donkey Anti-Chicken, 1:1000, Jackson Immunoresearch #703-545-155) was applied first to the slides for 90 min at room temperature. The secondary was washed off using 3×10 minute washes with PBT. Next, the mCherry secondary (Alexafluor 594 AffiniPure species-specific, 1:1000, Jackson Immunoresearch) was repeated for the same time and conditions. Finally, 3×10 minute washes with PBT were used to clear the slides of any remaining secondary and the slides were briefly dip-rinsed in PBS to remove Triton. The slides were wiped dry along back, sides, and edges and mounted using DAPI Fluoromount-G (Southern Biotech, #0100-20). PDE11A4-filled structures were quantified by an experimenter blind to treatment with images captured using Leica Application Suite (LASX) software and a Leica DM5000 B florescent microscope.

### Social Transmission of Food Preference

As previously described [22, 23], subjects’ access to food was restricted the two days prior to testing to one hour per day. The day prior to testing, all mice were placed in a clean home cage and given access to plain powdered chow packed into a glass jar. The following day, the designated “demonstrator mouse” was individually placed in a clean home cage and fed powdered chow flavored with a household spice (e.g. 5% orange vs 0.5% anise for the 24-hour test or 2% basil vs 1.5% thyme for the 7 day test). After one hour, the “demonstrator mouse” was returned to the original home cage where the “observer” cage mates were allowed unrestricted access to the demonstrator for 15 minutes. It is during this time that the observers make an association between the social pheromones in the breath of the demonstrator and the non-social odor (household spice). Recent and remote long-term memory were assessed 24 hours or 7 days after training, respectively. At that time, the observer mice were individually placed in clean home cages and given access to two flavored/powdered chows for 1 hour. One flavored chow contained a novel spice and the other contained the spice that their demonstrator was given. The amount of food eaten was measured by an experimenter blind to treatment. All mice met the minimum inclusion criteria of eating at least 0.25 grams of food. Note, the same cohort of mice was used to test both recent and remote memory, to reduce the total number of mice used since we have shown that this does not confound interpretation of the data [23]. Mice were allowed to recover from food restriction for ∼3 weeks prior to training for the second retrieval time point. Observer mice eating more food containing the familiar spice (i.e., the spice on their demonstrator’s breath) versus the novel spiced food constituted memory (preference ratio: familiar-novel/familiar+novel).

### Odor Recognition

As previously described [22, 23], subjects were allowed to habituate to 1’’ round wooden beads (Woodworks) for at least seven days prior to testing by placing several beads in the subjects’ home cages. For social odor recognition (SOR), the wooden beads are placed in home cages of other mice so the beads can take up the scent of different mouse strains (e.g., C57BL/6J Jax #000664, BALB/cJ Jax #000651, 129S6/ SvEv Taconic #129SVE). For non-social odor recognition (NSOR), the wooden beads are placed in a bag containing bedding saturated with a household spice (e.g., marjoram, cumin, etc.) for at least 7 days. Training for SOR and NSOR consisted of a habituation trial with 3 beads from the subject’s home cage, followed by two training trials that included 2 home-cage beads and 1 novel-scented bead. Recent and remote long-term memory were assessed 24 hours or 7 days after training, respectively. During SOR, mice were tested with one home cage bead, one bead from the trained donor strain (familiar), and one bead from a second donor strain (novel). During NSOR, mice were tested with only two beads, one scented with the training spice and one a novel spice. The designation of which scent was “novel” within a given testing trial and the location of the novel scent (i.e., left versus right) was counterbalanced across subjects. Mice were given two minutes to investigate the beads and the amount of time spent on each was manually scored by an experimenter blind to treatment and bead. For reasons not yet understood, we previously determined that infusion of even a negative control lentivirus into CA1 of hippocampus reverses the recent long-term memory impairment observed in *Pde11a* KO mice 24 hours after training [23]. Therefore, we did not test 24-hour SOR memory following injection of the isolated GAF-B domain as the results would not be interpretable. All mice met minimum inclusion criteria of spending a minimum of 3 seconds in total sniffing the beads. Spending more time investigating the novel vs familiar scent constituted memory (preference ratio: novel-familiar/novel+familiar).

### PDE and cyclic nucleotide assays

cAMP- and cGMP-PDE catalytic activity were measured as previously described [3]. The assay was validated *in vitro* using HT-22 cells (a mouse hippocampal cell line) as previously described [3]. Buffer containing 20 mM Tris-HCl and 10 mM MgCl_2_ was used to harvest cells and kept on ice until ready to use. PDE activity was measured using 50 µl of sample and 50 uL of of [_3_H] cAMP (Perkin Elmer, NET275) or cGMP (Perkin Elmer, NET337) and incubated for 10 minutes. After incubation, 0.1M HCL was added to quench the reaction, followed by 0.1M Tris to neutralize the reaction. 3.75 mg/mL snake venom (Crotalus atrox, Sigma V-7000) was then added to complete the reaction and the mixture was incubated at 37 °C for 10 minutes. Samples were put into 5’polystyrene chromatography columns with coarse filters (Evergreen, 208-3383-060) containing DEAE Sephadex A-25 resin (VWR, 95055-928). The columns were equilibrated in high salt buffer (20mM Tris-HCL, 0.1% sodium azide, and 0.5M NaCl) and low salt buffer (20mM Tris-HCL and 0.1% sodium azide). The reactions were then run down the equilibrated columns. Following four washes with 0.5 ml of low salt buffer, 4ml of Ultima Gold XR scintillation fluid (Fisher, 50-905-0519) was added to the eluate and mixed thoroughly. A Beckman-Coulter liquid scintillation counter Beckman LS 6000) was used to read counts per minute (CPM). As an assay control, two reactions free of sample lysate were ran in parallel to account for background activity and could then be subtracted from the sample CPMs. Total protein levels were quantified using the DC Protein Assay Kit (Bio-Rad, Hercules, CA) as described above, and CPMs were then normalized to the total amount of protein in each sample. To measure cAMP and cGMP levels directly, Cayman Chemical ELISA kits were used as per manufacturer’s instructions, as previously described [23].

### Biochemical Fractionation and Western Blotting

As previously described [1, 9, 26], biochemical fractionation was performed to obtain cytosolic and soluble membrane fractions. Cell were mixed with ice cold fractionation buffer (FB: 20 mM Tris-HCl, pH 7.5; 2 mM MgCl2; Thermo Pierce Scientific phosphatase tablet #A32959 and protease inhibitor 3 #P0044) and sonicated. First, a low-speed spin (1000 x g) removed cellular debris and the supernatant from this spin was transferred to a new tube. Next, a high-speed spin (89,000 x g) was performed to obtain the membrane (pellet) and cytosolic (supernatant) proteins. The pellet was rinsed and resuspended by sonication in fractionation buffer with 0.5% Triton-X 100. To solubilize the protein, samples were nutated for at 4°C for 30 minutes. A second high-speed spin (60,000 x g) was done for 30 minutes to separate the soluble membrane (supernatant) from the insoluble membrane (pellet). The soluble membrane sample was then transferred to a clean tube and used for western blot (see below). A DC Protein Assay kit (Bio-Rad; Hercules, CA, USA) was used to determine protein concentrations by which total protein was equalized across samples at 1 µg/µl. Samples were stored at −80°C until used in Western blotting. For Western blotting, 10ug of protein was loaded onto 12% NuPAGE Bis-Tris gels (Invitrogen, Waltham MA) and run at 180 volts. After about one hour, proteinwas transferred onto 0.45um nitrocellulose membrane using 100 mA for two hours. Following transfer, tris-buffered saline with 0.1% tween20 (TBS-T) was used to wash membranes. Membranes were cut into multiple strips to probe for multiple antibodies if needed. The membranes were blocked using either 5% milk or Superblock (PBS) Blocking Buffer (ThermoFisher, Cat#37515). Primary antibody was applied overnight at 4°C for PDE11A (Fabgennix PD11-101 at 1:500). The following day, membranes were washed with TBS-T (4×10 minutes). Secondary antibody (Jackson Immunoresearch Anti-Rabbit, 111-035-144; 1:10 000), was applied at room temperature for one hour. Finally, the membrane was washed in TBS-T (3×15 minutes). Chemiluminescence (SuperSignal West Pico Chemilumiscent Substrate; ThermoScientific, Waltham MA) was captured using film and multiple exposures were taken to ensure densities were within the linear range of the film. ImageJ was used to quantify optical densities. As previously described (e.g., [9, 12, 26]), each blot was normalized to a control condition (e.g., WT) to account for any technical variables (film exposure, antibody signal-noise, variance in chemiluminescence, etc.).

### Data Analysis

Data was collected by investigators blind to treatment and experiments were designed to counterbalance technical/biological variables. Outliers more than 2 standard deviations away from the mean were removed prior to analyses, as previously described (e.g., [9, 28, 29]). Outliers removed/total n: Figure 1A, 1/48; Figure 1J, 2/46; Figure 2A, 2/18; Figure 2B, 1/22; Figure 2D, 5/48; Figure 2E, 4/49; Figure 3A, 1/18; Figure 3B, 1/18; Figure 3C, 3/45;; Figure 3E, 2/20; Figure 3G, 9/112; Figure 3H, 2/35; Figure 3I, 3/48. Data were analyzed for effect of genotype, behavioral measure (e.g., bead or food), and treatment. Parametric statistical analyses were run on SigmaPlot 14.5 (San Jose, CA, USA) including ANOVA (F), Student’s t-test (t), and one-tailed one-sample t-test (t) when datasets met assumptions of normality (Shapiro-Wilk test) and equal variance (Brown-Forsythe test). To offset the possibility of a Type I error associated with multiple comparisons, a false-rate discovery (FDR) correction was applied to P-values from one-sample t-tests within an experiment, as previously described [3, 23]. If analyses failed normality and/or equal variance, nonparametric Kruskal-Wallis ANOVA (H) or Mann-Whitney rank sum test (T) were used instead. Student-Newman-Keuls or Dunn’s test were performed for *Post hoc* analyses. Significance was defined as P<0.05.

**Figure 1.**
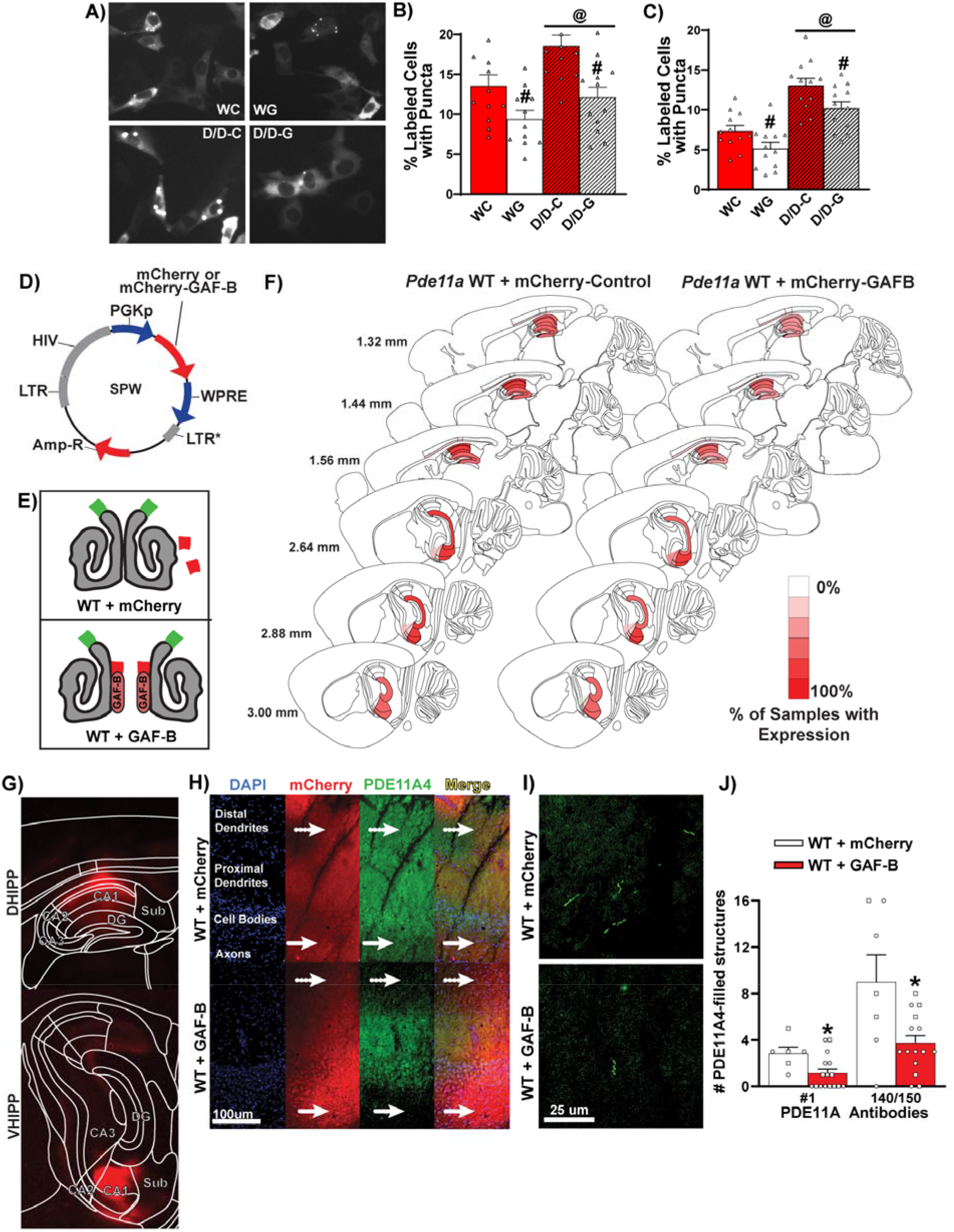
Disrupting PDE11A homodimerization in vivo selectively decreases PDE11A4 expression in a compartment-specific manner and is sufficient to reverse PDE11A4 accumulations in ghost axons that occur with age. PDE11A4-pS117/pS124 is a key intramolecular mechanism that drives the accumulation of PDE11A4 in ghost axons [2], so we conducted *in vitro* experiments to determine if an isolated GAF-B domain could reverse the effects of this phosphorylation. Indeed, disrupting homodimerization not only reduced the accumulation of WT PDE11A4 as previously described [1], it also reduced back to WT levels the potentiated accumulation caused by the phosphomimic mutant S117D/S124D (D/D) in both A,B) HT-22 cells (n=12 biological replicates/group; effect of S117D/S124D: F(1, 43)=8.80, P=0.005; effect of GAF-B: F(1,43)=16.74, P<0.001) and C) HEK293T (n=12 biological replicates/group; effect of S117D/S124D: F(1, 44)=46.54, P<0.001; effect of GAF-B: F(1,44)=10.31, P=0.002). D) A lentiviral construct containing either mCherry (i.e., negative control) or an mCherry-tagged isolated GAF-B domain (GAF-B) was injected bilaterally into dorsal and ventral CA1 of hippocampus. E) Previously, we showed the isolated GAF-B domain disrupts PDE11A4 homodimerization; whereas, mCherry alone does not [1]. F,G) Stereotaxic delivery of these lentiviral constructs to dorsal hippocampal (DHIPP) and ventral hippocampal (VHIPP) CA1 sub-regions resulted in high expression within CA1 in all subjects, with a subset of mice demonstrating a diffuse expression in dentate gyrus, CA2, and CA3. H) As previously reported [3], expression of mCherry-GAF-B decreased PDE11A4 expression in distal dendrites (dotted-arrows) and axons (solid arrows). I) The GAF-B construct was also able to reverse age-related increases in so-called “PDE11A ghost axons” (i.e., filamentous structures where PDE11A4 accumulates with age; shown: “140/150” cocktail of antibodies). J) In CA1 of old *Pde11a* wild-type (WT) mice, two different PDE11A4 antibody preparations revealed a significant decrease in PDE11A4-filled structures in GAF-B-treated mice (n=7M,9F) relative to mCherry-treated mice (n=3M,3F), including “#1” that detects all PDE11A4 (GAF-B, n=7M,9F; mCherry, n=3M,3F; fails normality, Rank Sum Test for effect of group: T(6,16) = 99.00, P=0.022) and a cocktail of “140/150” that specifically detects PDE11A4-pS117/pS124 (GAF-B, n=6M,9F; mCherry, n=3M,4F; fails normality, Rank Sum Test for effect of group: T(7,15) = 109.00, P=0.043). Brightness, histogram stretch, and/or contrast of images adjusted for graphical clarity.

**Figure 2.**
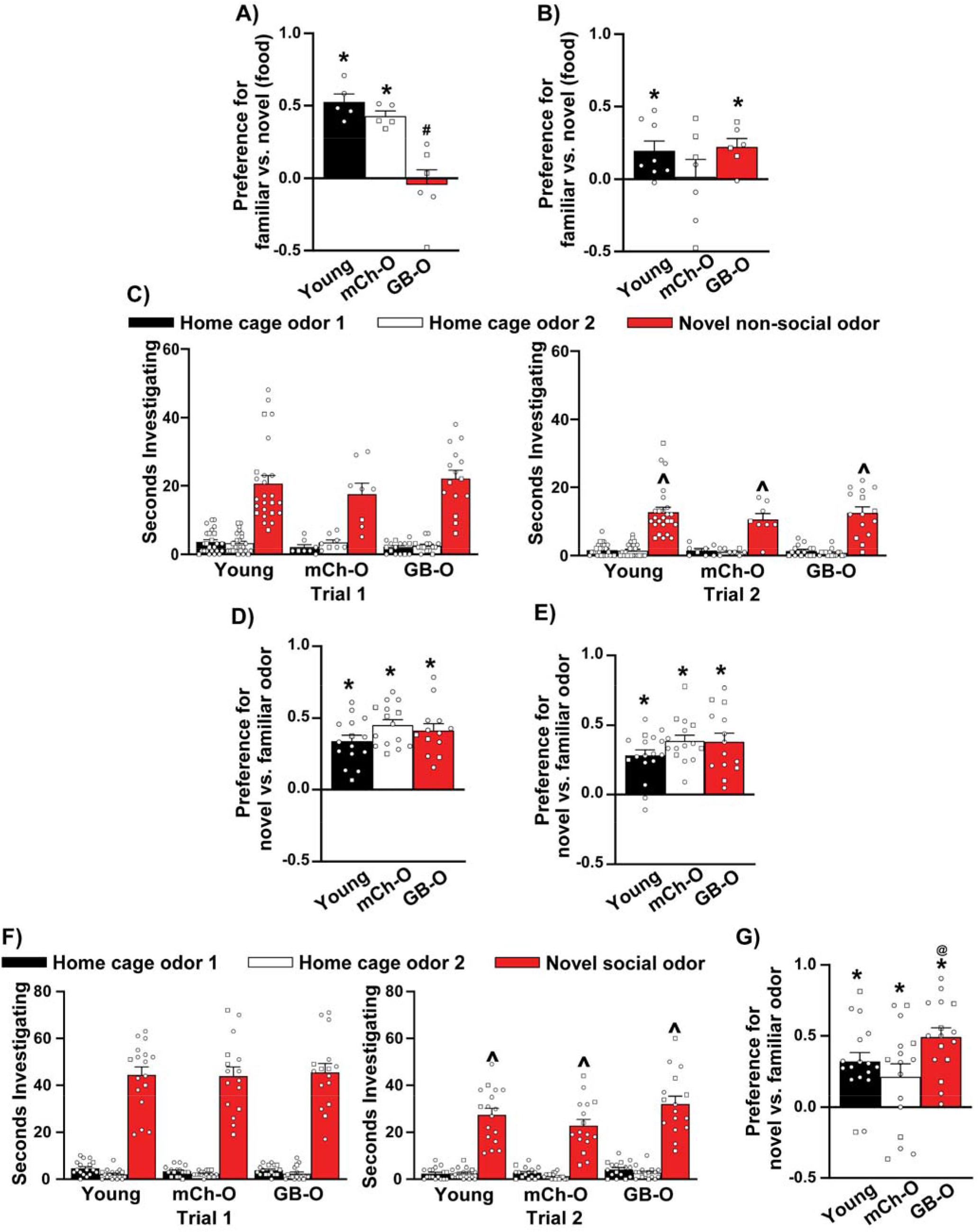
Disrupting PDE11A homodimerization in the hippocampus of old mice is sufficient to reverse age-related decline of remote long-term social memory. Previously, we determined that *Pde11a* WT mice suffer from age-related decline of remote long-term social associative memory (i.e., 7 days after training); whereas, genetic deletion of PDE11A prevents this ARCD, albeit at the expense of not being able to access ecent LTM 24 hours after training [2]. Therefore, we determined if disrupting homodimerization of PDE11A4 in CA1 of old *Pde11a* WT mice using the mCherry-tagged isolated GAF-B domain (GB-O) would be sufficient to produce such a transient amnesia that ultimately reverses ARCD of remote long-term social memory in old WT mice treated with mCherry alone (mCh-O). A) Indeed, using the social transmission of food preference assay (STFP), we found that GB-O mice exhibited impaired recent LTM relative to mCh-O and young mice (GB-O; n=3M/3F, mCh-O; n=3M/2F, Young; n= 5F; effect of group: F(2,13) = 16.52, P<0.001; *Post hoc*: GAF-B vs each group, P<0.001). B) In contrast, GB-O mice (n= 3M/3F) and young mice (n=8F) demonstrated significant remote LTM for STFP, but mCh-O mice did not (n= 4M/3F; Table 1). To determine if disrupting PDE11A4 homodimerization alters the non-social and/or social recognition components of the STFP assay, we tested odor recognition memory in these mice. C) GB-O and mCh-O mice learned equally well relative to young mice during non-social odor recognition (NSOR) training (GB-O, n=5M/10F; mCh-O, n=5M/11F; Young, n= 2M/15F; effect of trial for novel odor: F(1,45) = 76.58, P<0.001). D) All groups also demonstrated equivalent NSOR recent LTM (GB-O, n=3M/10F; mCh-O, n=4M/11F; Young, n= 2M/13F; effect of group: F(2,40) = 1.91, P=0.162) and E) NSOR remote LTM (GB-O, n=4M/11F; mCh-O, n=5M/10F; Young, n= 2M/13F; effect of group: F(2,42) = 0.53, P=0.590). F) Similarly, these GB-O (n=5M/11F) and mCh-O mice (n=5M/11F) learned equally well relative to young mice (n=2M/15F) during social odor recognition training (SOR) (effect of trial for novel odor: F(1,46) = 115.79, P<0.001). As previously reported in both young adult and old *Pde11a* KO mice [2], however, G) GB-O mice showed significantly improved remote SOR memory relative to mCh-O mice (GB-O, n=5M/10F; mCh-O, n=5M/11F; Young, n= 2M/15F; effect of group: F(2,45) = 3.76, P=0.031; *Post hoc*: mCh-O vs. GB-O P= 0.026). #vs Young, P=0.001; ^vs Trial 1, P<0.001; *has memory (i.e., significantly >0), P=0.023 to <0.001 (see Table 1 for one-sample t-tests); @vs mCh, P=0.026. Data plotted as individual points (females as circles, males as squares) and expressed as mean ±SEM.

**Figure 3.**
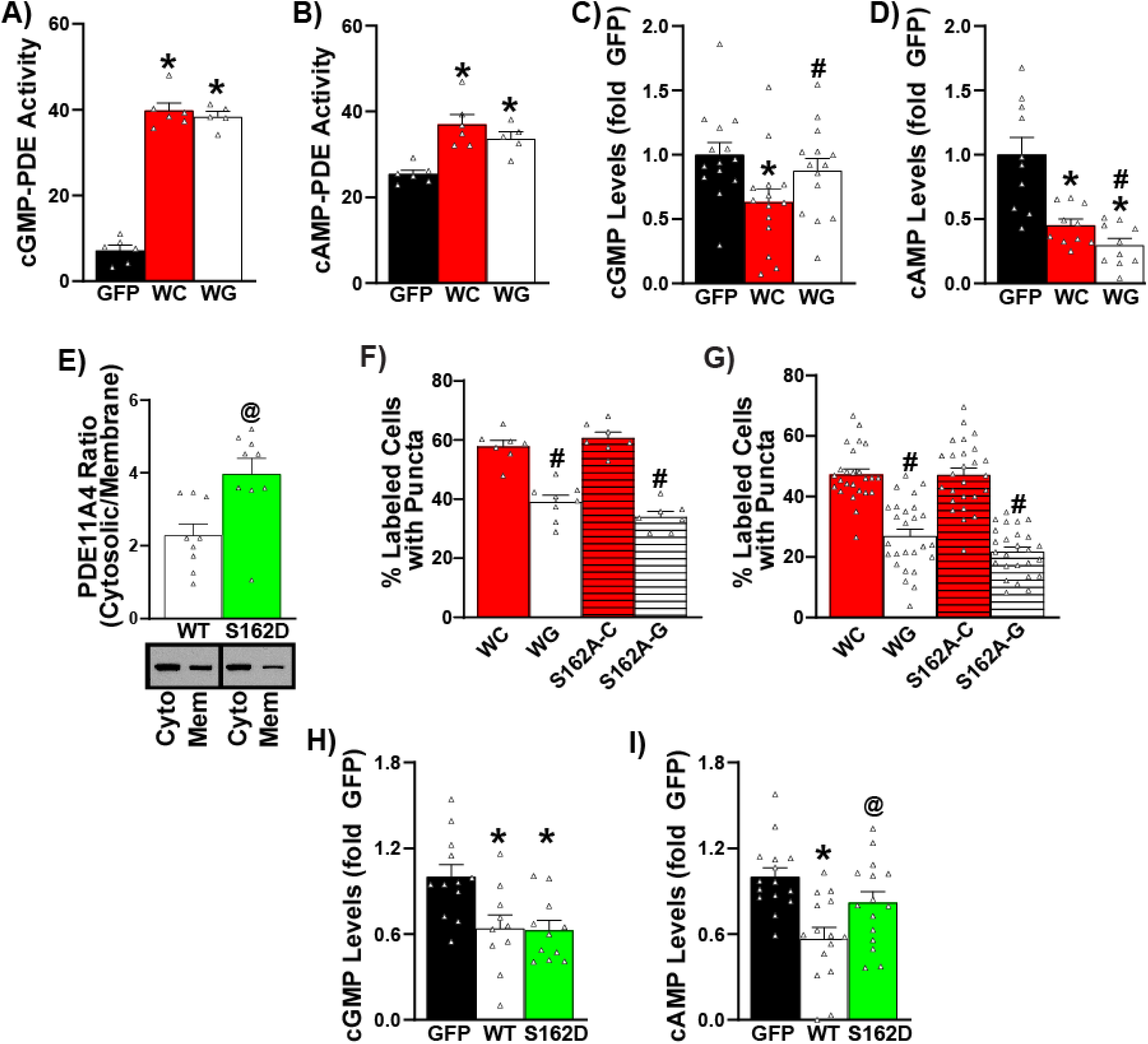
PDE11A homodimerization reverses biochemical phenotypes associated with aging independently of pS162 phosphorylation. Relative to transfection of GFP alone + mCherry alone (GFP, n = 6 biological replicates), HT-22 hippocampal cells transfected with either GFP-PDE11A-WT + mCherry (WC; n – 6 biological replicates) or GFP-PDE11A-WT + mCherry-GAF-B (WG; n=5 biological replicates) showed equivalent increases in both A) cGMP-PDE activity (effect of group: F(2,14)=168.01, P<0.001; *Post hoc*: GFP vs. all groups, P<0.001) and B) cAMP-PDE activity (effect of group: F(2,14)=12.85, P<0.001; *Post hoc*: GFP vs. WC P<0.001, GFP vs. WG P=0.005). Despite the fact that disrupting homodimerization did not directly affect PDE11A4 catalytic activity, it C) attenuated PDE11A4-induced decreases in cGMP levels (n=14 biological replicates/group; effect of group: F(2,39)=4.33, P=0.020; *Post hoc*: GFP vs. WC P=0.020, WG vs. WC P=0.041) while D) exacerbating PDE11A4-induced decreases in cAMP in COS1 cells(n=10 biological replicates/group; failed equal variance; ANOVA on Ranks for effect of group: H(2)=16.34, P<0.001; *Post Hoc*: GFP vs. all groups P<0.001, WC vs. WG P=0.034). The ability of the GAF-B construct to alter cyclic nucleotide levels in absence of a direct effect on PDE11A4 enzymatic activity is likely due to changes in subcellular compartmentalization of PDE11A4 that are described above [1]— possibly shifting it from a cGMP-rich pool to a cAMP-rich pool. Since the ability of the isolated GAF-B domain to disperse PDE11A4 accumulations strongly resembles that of a phosphomimic mutation at S162 (i.e., S162D [2]), we sought to determine if disrupting homodimerization elicited its effects by promoting phosphorylation of S162. E) As previously reported for the GAF-B construct [1], S162D shifts PDE11A4 from the membrane to the cytosol relative to WT (n=9 biological replicates/group; effect of group: t(16)=-3.21, P=0.005). F) That said, GAF-B is still able to reduce the accumulation of PDE11A4 in HEK293T cells even when a phosphomutant alanine is introduced at S162 (i.e., S162A; n=12 biological replicates/group; effect of GAF-B: F(1,26)=135.65, P<0.001). G) This effect was also observed in COS1 cells (n=25 biological replicates for each WT group, n=26 biological replicates for each S162A group; effect of GAF-B: F(1,99)=137.52, P<0.001). This suggests that disrupting homodimerization does not achieve effects by promoting phosphorylation of S162. Indeed, S162D differed substantially from GAF-B in terms of regulating cyclic nucleotide levels, with H) no effect on PDE11A4-induced decreases in cGMP levels (n=10-12 biological replicates/group; effect of group: F(2,30)=6.99, P=0.003; *Post hoc* GFP vs. WT P=0.004, GFP vs. 162D P=0.007, WT vs 162D P=0.924) and an I) an attenuation of PDE11A4-induced decreases in cAMP levels (n=15 biological replicates/group; effect of group: F(2,42)=8.76, P<0.001; *Post hoc* GFP vs. WT P<0.001, WT vs. 162D P=0.020, GFP vs 162D P=0.089). *vs. GFP P=0.02 to P<0.001; #vs. mCherry P=0.041 to P<0.001; @vs. WT, P=0.02 to P<0.001. Data plotted as individual data points and mean ±SEM.

## RESULTS

### Disrupting PDE11A4 homodimerization in the aged hippocampus selectively decreases PDE11A4 expression and accumulation in a compartment-specific manner

We previously found that ventral hippocampal PDE11A4 expression increases with age in both mice and humans and that these age-related increases accumulate specifically in the membrane compartment and within filamentous structures termed “ghost axons” due to increased phosphorylation of PDE11A4 at S117 and S124 [2]. Additionally, we found *in vitro* that disrupting PDE11A4 homodimerization by expressing an isolated PDE11A4 GAF-B domain (Figure 1B) leads to proteolytic degradation specifically of membrane-bound PDE11A4 and reduces the punctate accumulation of PDE11A4 [1]. Therefore, we sought to determine if disrupting PDE11A homodimerization would be sufficient to reverse age-related PDE11A4 proteinopathies in a compartment-specific manner. To do this, we first determined if disrupting PDE11A4 homodimerization would prevent the punctate accumulation not only of PDE11A4-WT (as previously described [1]), but also of PDE11A4-S117D/S124D, which mimics the age-related increase in PDE11A4-pS117/pS124 that drives the punctate accumulation of PDE11A4 in the aged brain [2]. Indeed, across multiple experiments and cell lines (i.e., HT-22 and HEK293T), the isolated GAF-B domain reduced the punctate accumulation of both PDE11A4-WT and PDE11A4-S117D/S124D compared to the mCherry control (Figure 1A-C). Next, we used lentiviruses (Figure 1D) containing either mCherry alone (i.e., negative control) or an mCherry-tagged isolated GAF-B domain that disrupts PDE11A4 homodimerization (Figure 1E; [1]). These lentiviruses were stereotaxically injected bilaterally into the CA1 field of dorsal and ventral hippocampus of old *Pde11a* WT mice, since this is the field where PDE11A4 regulates social learning and memory [23] (Figure 1F). In select experiments, young mice were also injected with the mCherry-only virus. All mice demonstrated mCherry signal in dorsal and ventral CA1, with a subset of mice exhibiting expression in neighboring hippocampal sub-regions (e.g., dentate gyrus, CA3, CA2) that do not express PDE11A4 (Figure 1E). While mCherry-treated mice exhibit a uniform pattern of PDE11A4 expression across stratum radiatum, stratum pyramidale and stratum oriens of CA1, GAF-B treated mice exhibit a significant compartment-specific decrease in PDE11A expression, with reduced expression in the distal segment of stratum radiatum and stratum oriens relative to stratum pyramidale (see [3] for quantification of effect; Figure1H). Consistent with the fact that the isolated GAF-B domain reduces PDE11A4 protein expression in the membrane compartment [1], we found that disrupting homodimerization of PDE11A4 reduced age-related increases in so-called “PDE11A4 ghost axons” (i.e., filamentous structures harboring age-related accumulations of PDE11A4 [2]; Figure 1I). The ability of the GAF-B construct to reduce age-related accumulations of PDE11A4 in ghost axons was confirmed using two different PDE11A4 antibodies (one that detects all PDE11A4 and another that detects PDE11A4-pS117/pS124 specifically) on two different sets of slides from the same cohort of mice (Figure 1J). All together, these data suggest disruption of PDE11A4 homodimerization *in vivo* is sufficient to reverse age-related increases in PDE11A4 protein expression and ectopic accumulation.

### Disrupting PDE11A4 homodimerization in the hippocampus of old mice is sufficient to reverse age-related decline of remote long-term social memory

Previously, we found that while *Pde11a* WT mice suffer from ARCD of remote long-term social associative memories, *Pde11a* KO mice do not [2]. Interestingly, this protection of remote LTM 7 days after training comes at the expense of not being able to access recent LTM 24 hours after training [2]. Therefore, we determined if disrupting homodimerization of PDE11A4 using the isolated GAF-B domain would be sufficient to induce a transient amnesia in old *Pde11a* WT mice that ultimately reverses ARCD of remote long-term social associative memories. Here we measure social associative memory using social transmission of food preference (STFP), an assay where mice form an association between a non-social odor (i.e., a household spice) and a social odor (i.e., pheromones in their cage mate’s breath), with the memory of that association indicating a food with that scent is safe to eat [30, 31]. Indeed, mice treated with the GAF-B domain showed no recent LTM for STFP (Figure 2A; Table 1) but did show remote LTM for STFP on par with that of young adult mice (Figure 2B; Table 1); whereas, mice treated with mCherry alone showed ARCD of remote LTM for STFP (Figure 2B; Table 1). Importantly, we found no significant differences between groups in terms of the total amount of food eaten (Table 2). To disentangle effects of the GAF-B construct on the non-social versus social components of the STFP assay, we turned to memory tests for odor recognition. Old *Pde11a* WT mice treated with mCherry alone or mCherry-GAF-B spent the same amount of time sniffing the beads (Table 2) and learned equally well during training for non-social odor recognition (NSOR; Figure 2C). They also demonstrated equally strong recent and remote LTM for NSOR (Figure 2D-E; Table 1). This suggests that disrupting PDE11A4 homodimerization— like genetically deleting PDE11A [22, 23]—does not alter the ability to detect, learn about, or retrieve memories for recognizing non-social odors [22]. In contrast, disrupting PDE11A4 homodimerization—again, like genetically deleting PDE11A [22, 23]—did significantly improve remote LTM for SOR memory (Figure 2G; Table 1), despite having no effect on SOR learning (Figure 2F) or total time sniffing (Table 2). These data suggest that disrupting PDE11A4 homodimerization is sufficient to reverse ARCD of remote social memory.

**Table 1.** Preference ratios for each group calculated and subequently analyzed by one-sample t-test to determine if their performance significantly differed from chance (i.e., 0) to determine whether or not a given group exhibited a signifciant memory. The resulting P-values within an experiment were then corrected for multiple comparisons using false discovery rate correction (FDR).

| Datasets | Young | mCh-O | GB-O |
| --- | --- | --- | --- |
| **Fig2A STFP 24h LTM** | t(4)=9.51, FDR-P<0.001 | t(4)=12.60, FDR-P<0.001 | t(6)=-0.043, FDR-P=0.344 |
| **Fig2B STFP 7d LTM** | t(7)=2.90, FDR-P=0.017 | t(6)=0.14, FDR-P=0.896 | t(5)=3.84, FDR-P=0.018 |
| **Fig2D NSOR 24h LTM** | t(14)=8.19, FDR-P<0.001 | t(14)=11.9, FDR-P<0.001 | t(12)=8.48, P<0.001 |
| **Fig2E NSOR 7d LTM** | t(14)=9.75, FDR-P<0.001 | t(14)=9.45, FDR-P<0.001 | t(14)=8.04, FDR-P<0.001 |
| **Fig2G SOR 7d LTM** | t(16)=7.49, FDR-P<0.001 | t(15)=2.18, FDR-P=0.023 | t(14)=6.74, FDR-P<0.001 |

**Table 2.**
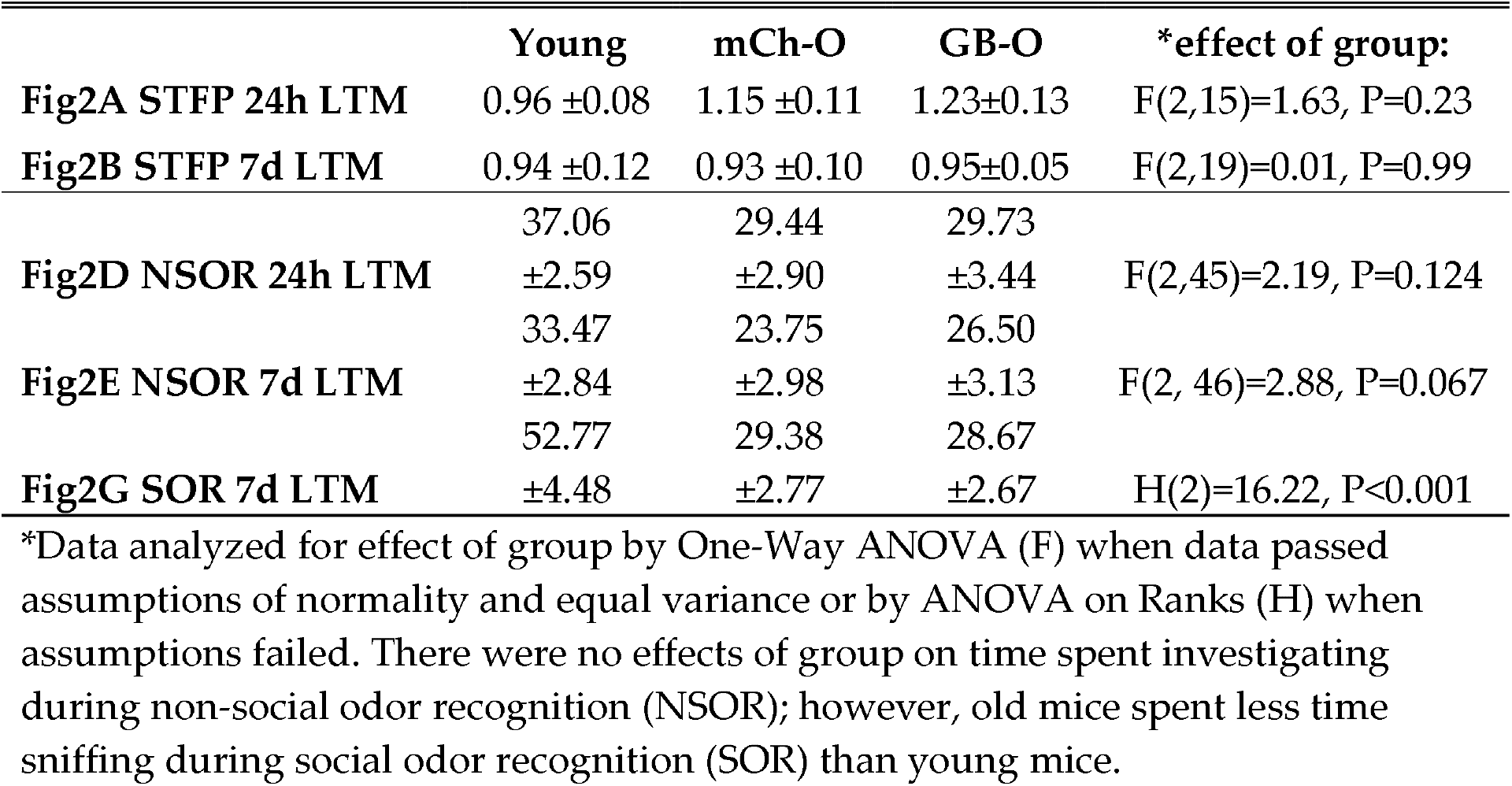
Amount of food (grams) eaten during STFP tests or amount of time spent investigating (seconds) during SOR/NSOR tests by young controls and old mice treated with mCherry (mCh-O) or GAF-B (GB-O) lentivirus during 24-hour and 7-day long-term memory (LTM) tests (Data expressed as mean ±SEM).

### PDE11A4 homodimerization is an independent intramolecular mechanism that regulates PDE11A4 trafficking and functioning

As noted above, disrupting PDE11A4 homodimerization significantly changes the subcellular compartmentalization of the enzyme. To better understand the functional consequences of disrupting PDE11A4 homodimerization, we measured PDE catalytic activity and cyclic nucleotide levels in cells transfected with GFP + mCherry (i.e., negative control), GFP-PDE11A4 + mCherry, or GFP-PDE11A4 + mCherry-GAF-B. Compared to the negative control, expression of PDE11A4 significantly increased cGMP- and cAMP-PDE activity in HT-22 cells as expected. This increase in PDE activity was not altered by disruption of PDE11A4 homodimerization (Figure 3A-B), which is consistent with previous studies using purified PDE11A4 enzyme [20]. Also as expected, expression of PDE11A4 in concert with mCherry decreased both cGMP and cAMP levels in COS1 cells relative to the negative control (Figure 3C-D). Interestingly, disrupting homodimerization of PDE11A4 decreased PDE11A4-mediated degradation of cGMP while exacerbating PDE11A4-mediated degradation of cAMP (Figure 3C-D). Together, these data suggest disruption of PDE11A4 homodimerization alters cyclic nucleotide signaling not by altering PDE11A4 catalytic activity directly, but rather by changing the subcellular localization of PDE11A4 protein (e.g., shifting from a cGMP-rich pool to a cAMP-rich pool).

### PDE11A homodimerization does not require phosphorylation of S162

As described above, disrupting PDE11A4 homodimerization reduces the accumulation of PDE11A4 in punctate structures both *in vivo* (Figure 1F) and *in vitro* ([1]; Figure 3E). Such a dispersing phenotype is also triggered *in vitro* by a phosphomimic mutation at S162 (i.e., S162D) [2]. As such, we sought to determine if disrupting PDE11A4 homodimerization may work by promoting phosphorylation of S162. Indeed, S162D changed the subcellular compartmentalization of PDE11A4 in a manner similar to that observed with the isolated GAF-B domain [1]—namely, shifting PDE11A4 from the membrane to the cytosolic fraction (Figure 3E). That said, the isolated GAF-B domain was able to effectively reduce the accumulation of both PDE11A4-WT as well as the phosphoresistant PDE11A4-S162A, suggesting phosphorylation of S162 is not needed for the dispersing effect of the isolated GAF-B domain (Figure 3IF-G). Consistent with this suggestion, we found that S162D elicited quite different effects on cyclic nucleotide levels than were described above for the isolated GAF-B domain. Specifically, S162D did not alter PDE11A4 hydrolysis of cGMP (Figure 3H) and reduced PDE11A4 hydrolysis of cAMP (Figure 3I). Together these data suggest that homodimerization and pS162 are independent intramolecular mechanism that regulate PDE11A4 trafficking and function.

## DISCUSSION

Previously, we found that age-related increases in PDE11A4 occur specifically within the membrane compartment of the VHIPP and ectopically accumulate in filamentous structures we term “ghost axons” (see [2]) due to age-related increases in the phosphorylation of PDE11A4-S117/S124. Here, we show that disrupting PDE11A4 homodimerization within hippocampal CA1 using a biologic encoding an isolated GAF-B domain reverses the accumulating effect of S117D/S124D *in vitro* (Figure 1A-C), reduces PDE11A4 expression in ventral CA1 in a compartment-specific manner (i.e., in distal dendrites and stratum oriens; Figure 1H), and reverses the age-related accumulation of PDE11A4 in ghost axons *in vivo* (Figure 1I-J). Further, we show that disrupting PDE11A homodimerization is sufficient to reverse ARCD of remote social aLTMs—albeit at the expense of an inability to retrieve recent social LTMs (Figure 2). Such a transient amnesia that ultimately produces stronger remote social LTMs is also observed with genetic deletion of *Pde11a* in young and old adult mice [2, 23]. Although disrupting PDE11A homodimerization does not alter catalytic activity of the enzyme, it ultimately leads to higher cGMP levels and slightly lower cAMP levels (Figure 3C-D) by virtue of changing the subcellular localization of PDE11A4 (Figure 3E-G). This suggests that disrupting PDE11A4 homodimerization shifts PDE11A from a cGMP-rich pool to a cAMP-rich pool, thereby alleviating age-related decreases in cGMP that are widely reported to occur in the aging hippocampus [6]. Interestingly, phosphorylation of PDE11A4 at serine 162 (pS162) similarly reduces the accumulation of PDE11A4 in punctate structures and shifts PDE11A4 from the membrane to the cytosol (Figure 3E) as does disruption of homodimerization [1, 2]. That said, we show here that these two regulatory mechanisms clearly operate independently since disrupting homodimerization 1) reduces the accumulation of PDE11A4 even when phosphorylation of S162 was prevented (Figure 3I-J) and 2) decreased cAMP levels (Figure 3D) while mimicking phosphorylation of S162 increased cAMP levels (Figure 3L). Together, these data suggest that disrupting PDE11A4 homodimerization *in vivo* via expression of its isolated GAF-B domain is sufficient to reverse the age-related ectopic accumulation of PDE11A4 within ghost axons and rescue ARCD of remote social aLTMs independently of phosphorylating S162.

### Post-translational modifications and protein-protein interactions alter the subcellular localization and trafficking of PDE11A4 protein

The subcellular compartmentalization of PDEs are regulated by multiple mechanisms [7]. The localization of the PDE2, PDE4, PDE5, PDE10, and PDE11 families is regulated in part by post-translational modifications (c.f., [7]). For instance, phosphorylation of PDE10A prevents membrane insertion by blocking palmitoylation [32]. As opposed to direct membrane insertion, PDE11A4 is thought to associate with the membrane by binding to a macromolecular complex [2]. The ability of PDE11A4 to interact with this macromolecular complex appears to be reduced when PDE11A4 homodimerization is disrupted [1] or when S162 is phosphorylated (Figure 3E), as both intramolecular signals cause PDE11A4 to shift from the membrane to the cytosol. Interestingly, this shift away from the membrane compartment following disruption of homodimerization or introduction of the S162D phosphomimic mutation is accompanied by a reduction in the punctate accumulation of PDE11A4 [1, 2]. Despite these similarities, however, the two regulatory mechanisms have opposing effects on cAMP levels (Figure 3D vs. 3K). These opposing effects suggest PDE11A4 is being redistributed to differing cytosolic compartments in response to disrupted homodimerization versus phosphorylation of S162. Indeed, disrupting homodimerization acts independently of pS162 to reduce the punctate accumulation of PDE11A4 (Figure 3H-I). Phosphorylation of PDE11A4 at serines 117 and 124 (pS117/pS124), on the other hand, increases expression and accumulation of PDE11A4 protein [2]. PKA and PKG have been shown to phosphorylate S117 and S162 *in vitro* [33], and these same kinases are predicted to phosphorylate S124 [19]. Cyclic nucleotide levels controlled by PDE11A4 could then modulate PKA/PKG activity at S117, S124, and/or S162, thereby creating direct feedback/feedforward loop as has been described for other PDE families [7].

In addition to post-translational modification, protein-protein interactions regulate the subcellular localization of PDEs [7]. Our work here (and elsewhere [1, 3]) suggests that homodimerization is a key type of protein-protein interaction that regulates the compartmentalization of PDE11A4 both *in vitro* and *in vivo*. For example, the PDE11A4 sequence in BALB/cJ mice differs from that of C57BL/6J mice at a single amino acid that falls within the GAF-B domain—whereas BALB/cJ mice encode a threonine at amino acid 499, C57BL/6J mice encode an alanine [1]. This A499T BALB/cJ mutation strengthens homodimerization, elevates protein expression, and increases the punctate accumulation of PDE11A4 both *in vitro* and *in vivo* [1, 3]. In contrast, disrupting homodimerization using the isolated GAF-B domain has the opposite effects, lowering protein expression of PDE11A4 due to increased proteolysis and reducing the punctate accumulation of PDE11A4 both *in vivo* and *in vitro* ([1, 3]; Figure 1E-G; Figure 3E-F and 3H-I). Homodimerization/oligomerization likely controls the subcellular compartmentalization of other GAF domain-containing PDE families, given GAF-B mutations change the subcellular compartmentalization of PDE6C [34]. The fact that disrupting PDE11A4 homodimerization lowers PDE11A4 expression and its punctate accumulation suggests that age-related increases in PDE11A4 expression/accumulation may be prevented/reversed by reducing levels of PDE11A4 homodimerization. Indeed, we show here that disrupting PDE11A4 homodimerization in old WT mice was sufficient to reverse age-related accumulation of PDE11A4 in ghost axons (Figure 1F-G) and ARCD of remote social aLTMs (Figure 2B).

### Disrupting PDE11A4 homodimerization as a therapeutic approach for treating ARCD

A hallmark pathology of the aging brain includes ectopic protein expression and accumulation in the brain [35], as well as the loss of cGMP signaling in the hippocampus [6, 36, 37]. Much like hyperphosphorylated tau causes neurofibrillary tangles that lead to impaired cell communication, function, and death [38], we find that phosphorylation of PDE11A4 at serines S117 and S124 increases PDE11A4 protein expression and ectopic accumulation in filamentous structures termed “ghost axons” [2]. These age-related increases in PDE11A protein appear to cause ARCD of social memories via the cGMP-PKG, as opposed to cAMP-PKA, pathway [2]. This is consistent with reports that aging and ARCD are associated with decreases in cGMP, but not cAMP, in the hippocampus [6, 36, 37], and that elevating cGMP levels elicits nootropic effects in the context of ARCD and neurodegenerative diseases [39-44]. Therefore, it is particularly noteworthy that disrupting PDE11A4 homodimerization increases cGMP levels (Figure 2), possibly by removing PDE11A4 from hydrolyzing a key nanodomain of cGMP.

Although it is clear that age-related increases in PDE11A4 protein expression are deleterious to remote social memory, it is important to consider the possibility that the ectopic accumulation of PDE11A in ghost axons may actually serve as a protective mechanism to sequester and/or neutralize excess PDE11A4 [2, 45]. An example of this type of sequestration is found with some PDE4A isoforms when they are bound by conformationally-altering catalytic inhibitors (e.g., [46]). Therefore, therapeutic targeting of PDE11A4 intended to reduce its ectopic accumulation may also need to promote clearance of PDE11A4 from the system. As noted above, disrupting PDE11A4 homodimerization significantly decreases the puctate accumulation of PDE11A4 and specifically promotes degradation of membrane-associated PDE11A4 [1]. Our results here suggest that disruption of PDE11A4 homodimerization may be a sophisticated mechanism for therapeutic targeting of age-related increases in PDE11A4, by decreasing both its accumulation and clearing it from select compartments. All together, these data suggest that disruption of PDE11A4 homodimerization using either a small molecule approach or a biologic—such as the biologic described herein—may represent a novel therapeutic approach for treating ARCD of social memories.

## Supporting information

Unprocessed images

## ACKNOWLEDGEMENTS

The authors would like to thank Dr. Mythreye Karthikeyan for mycoplasma testing of cell cultures as well as Anjali Pathak and Abigail Smith for technical assistance.

