## Supplementary figures and images for "Biologic that disrupts PDE11A4 homodimerization in hippocampus CA1 reverses age-related proteinopathies in PDE11A4 and cognitive decline of social memories"

### Unprocessed images

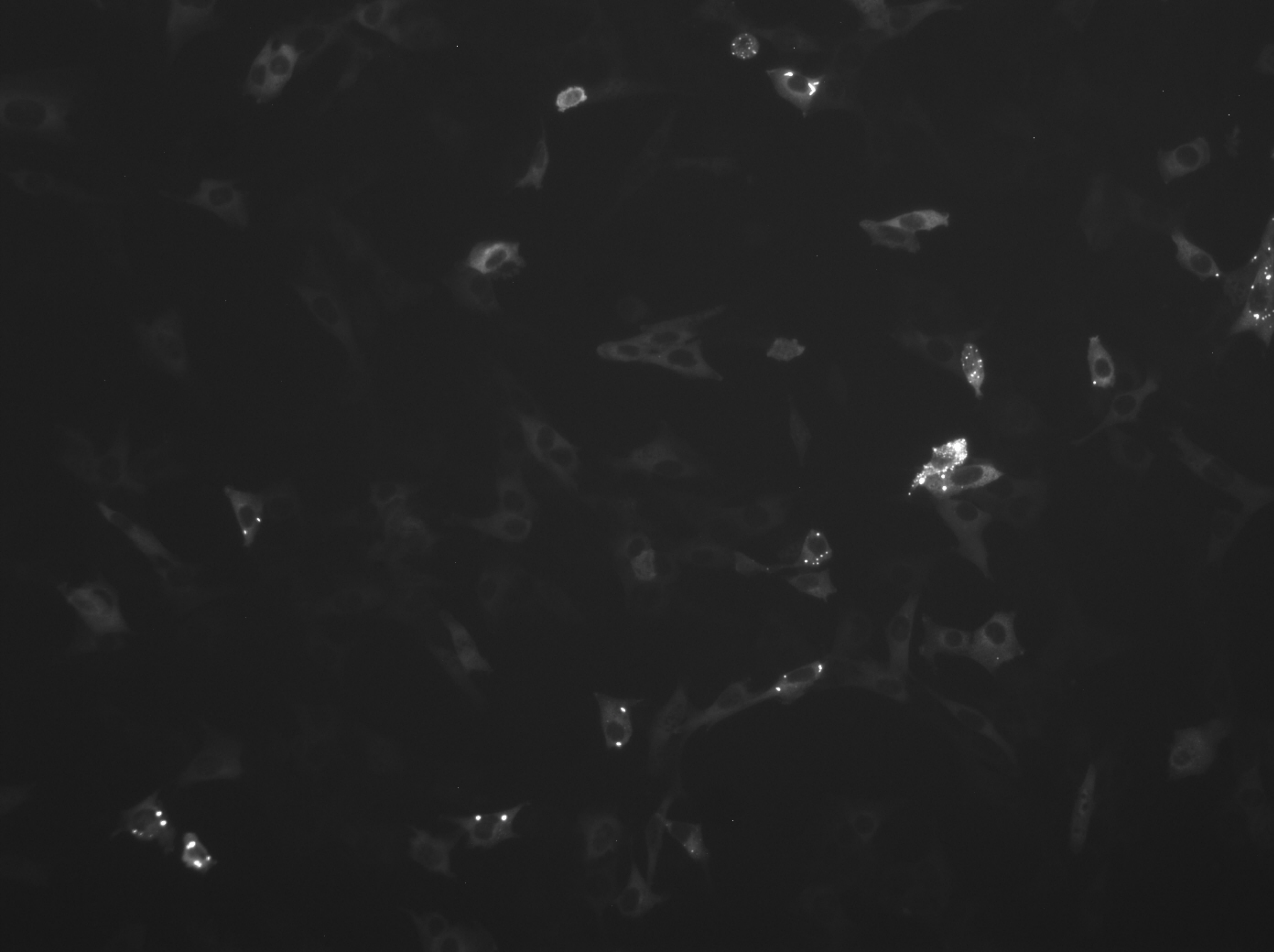

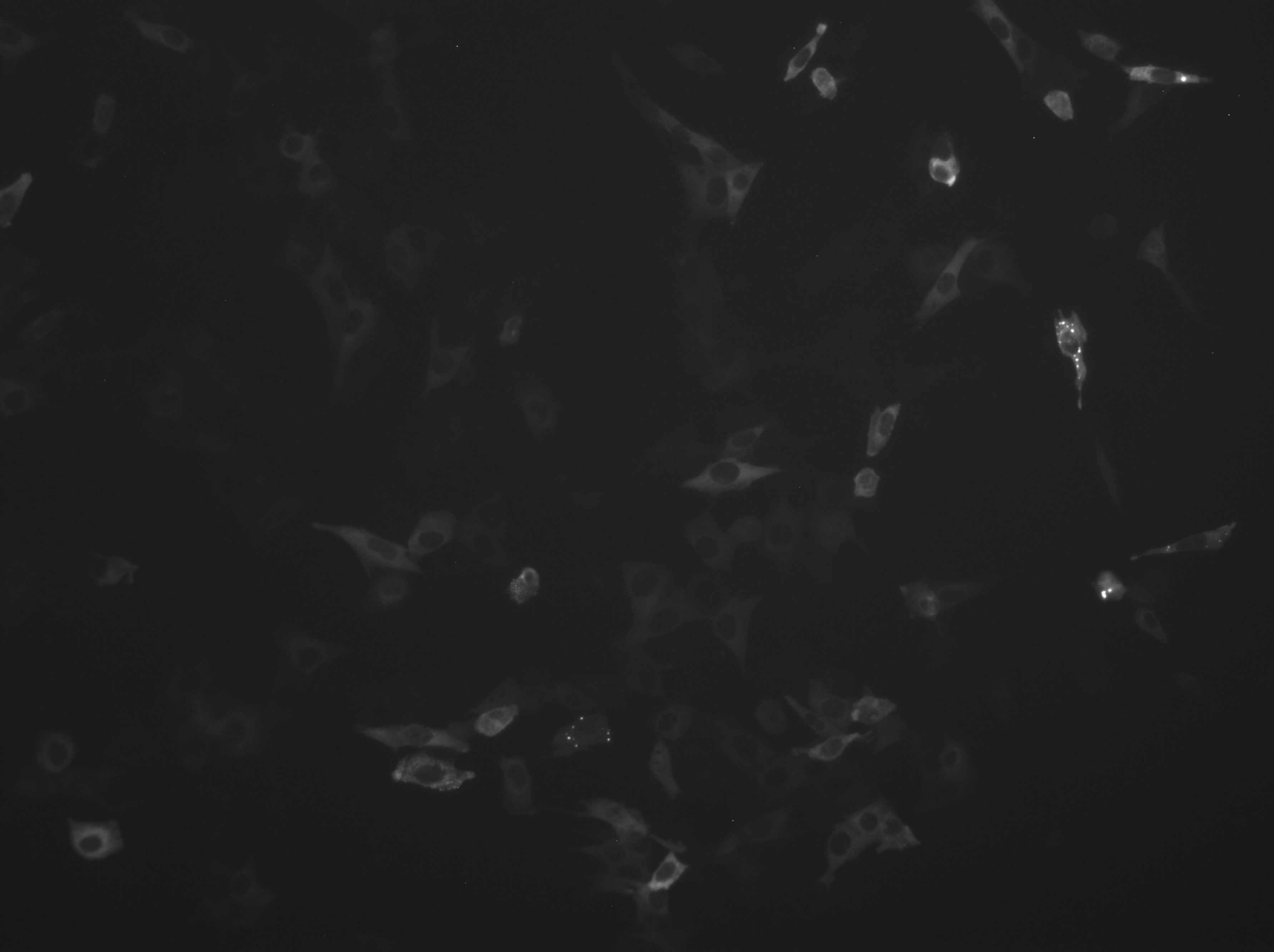

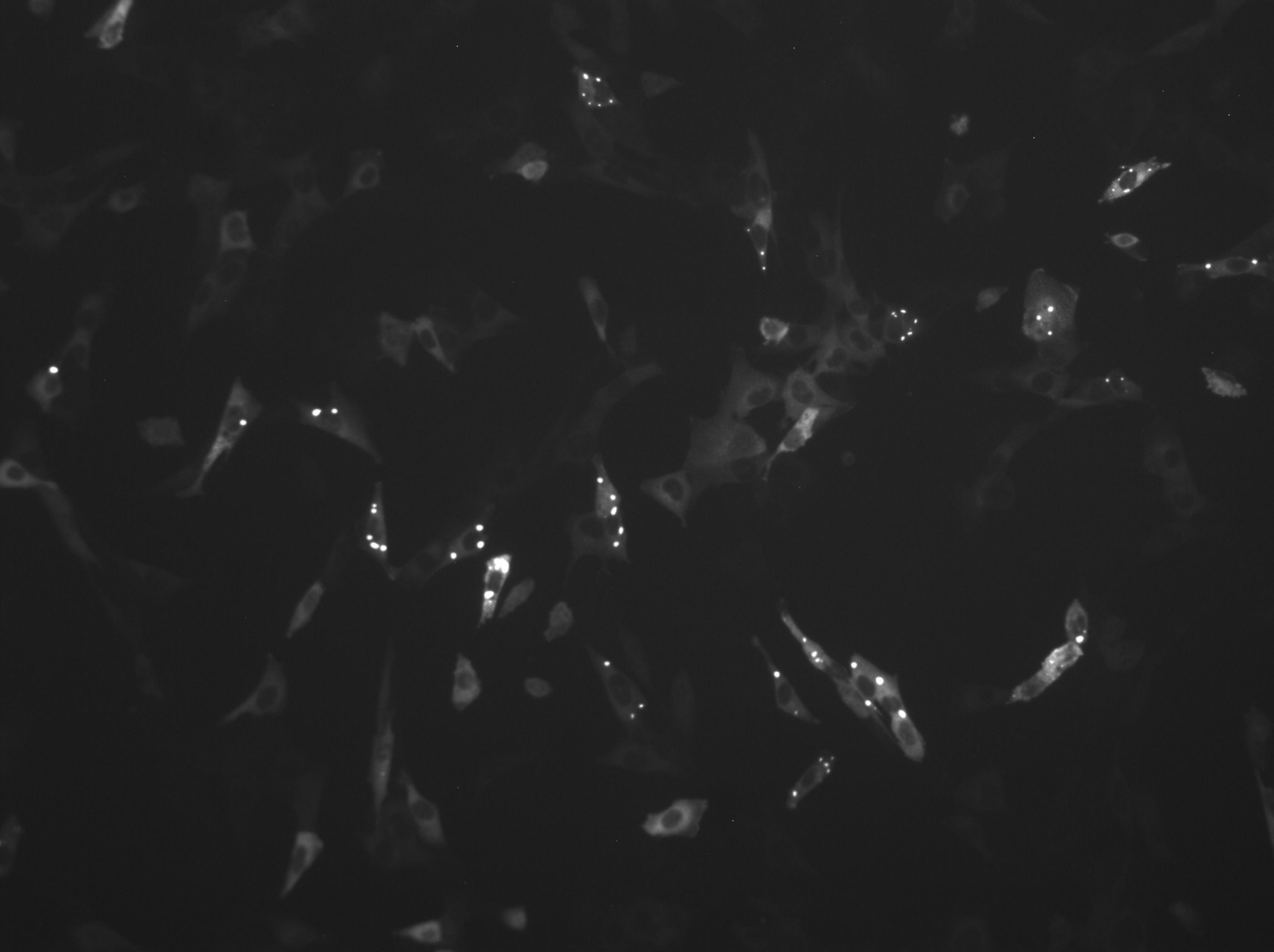

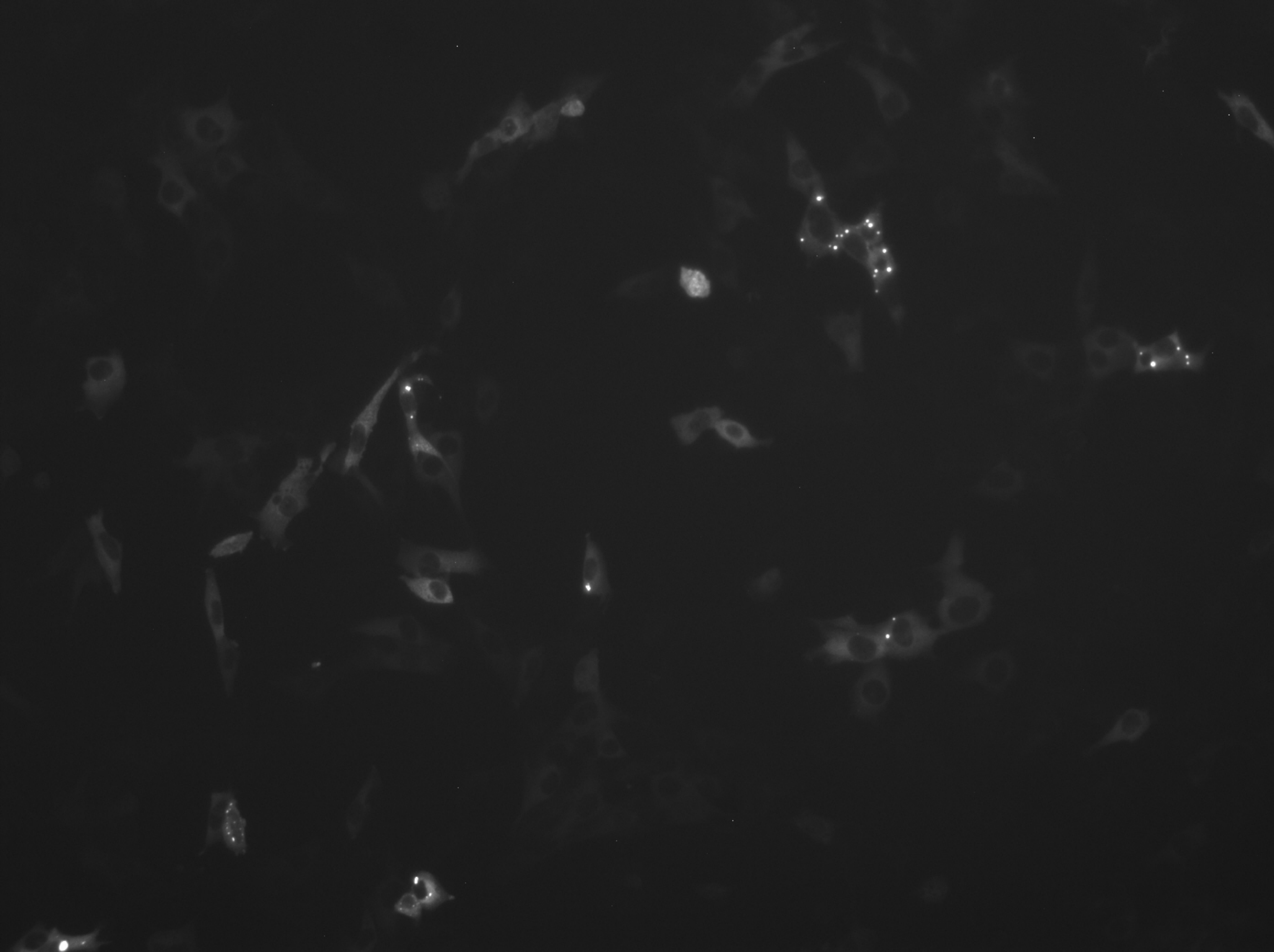

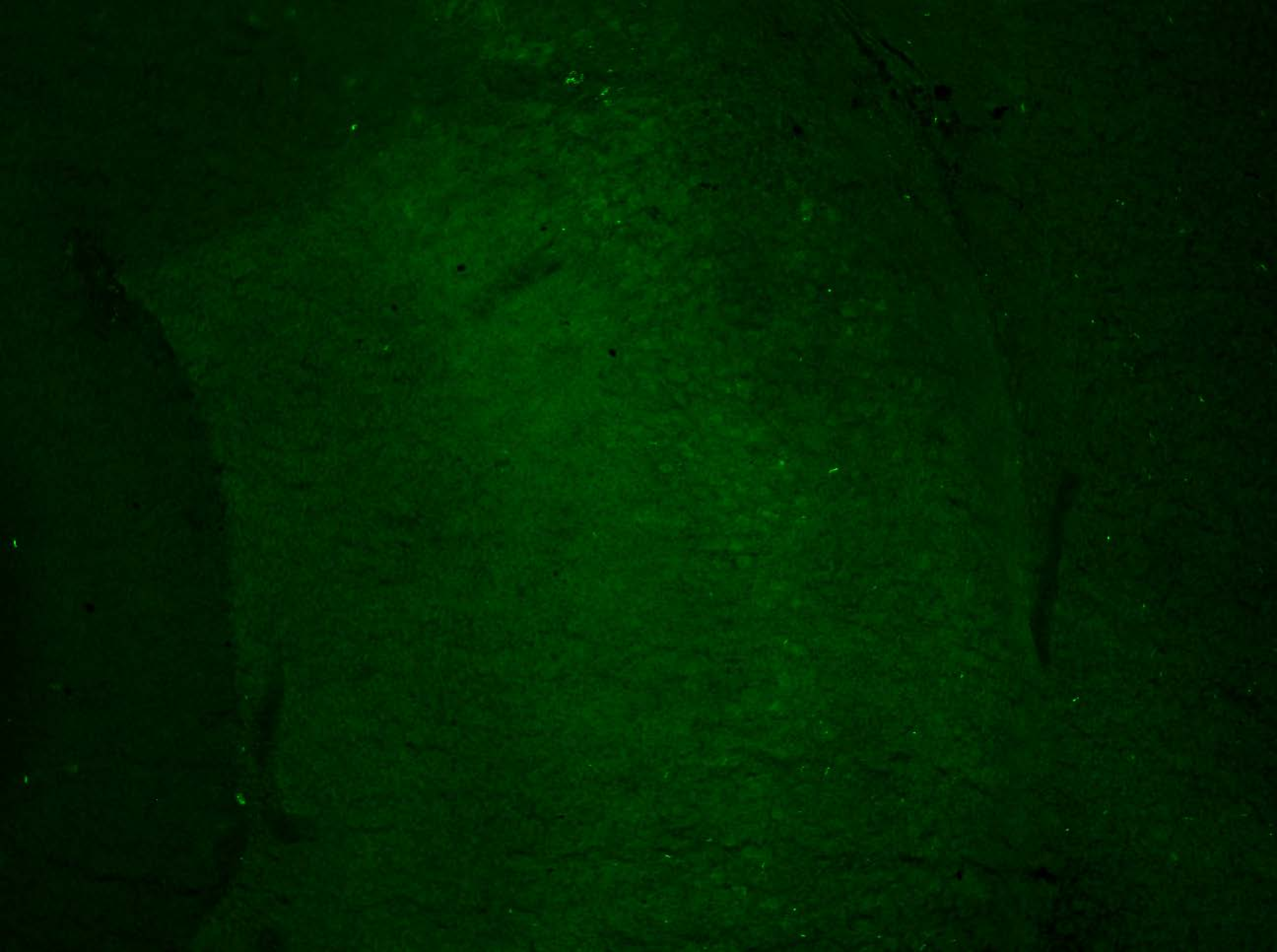

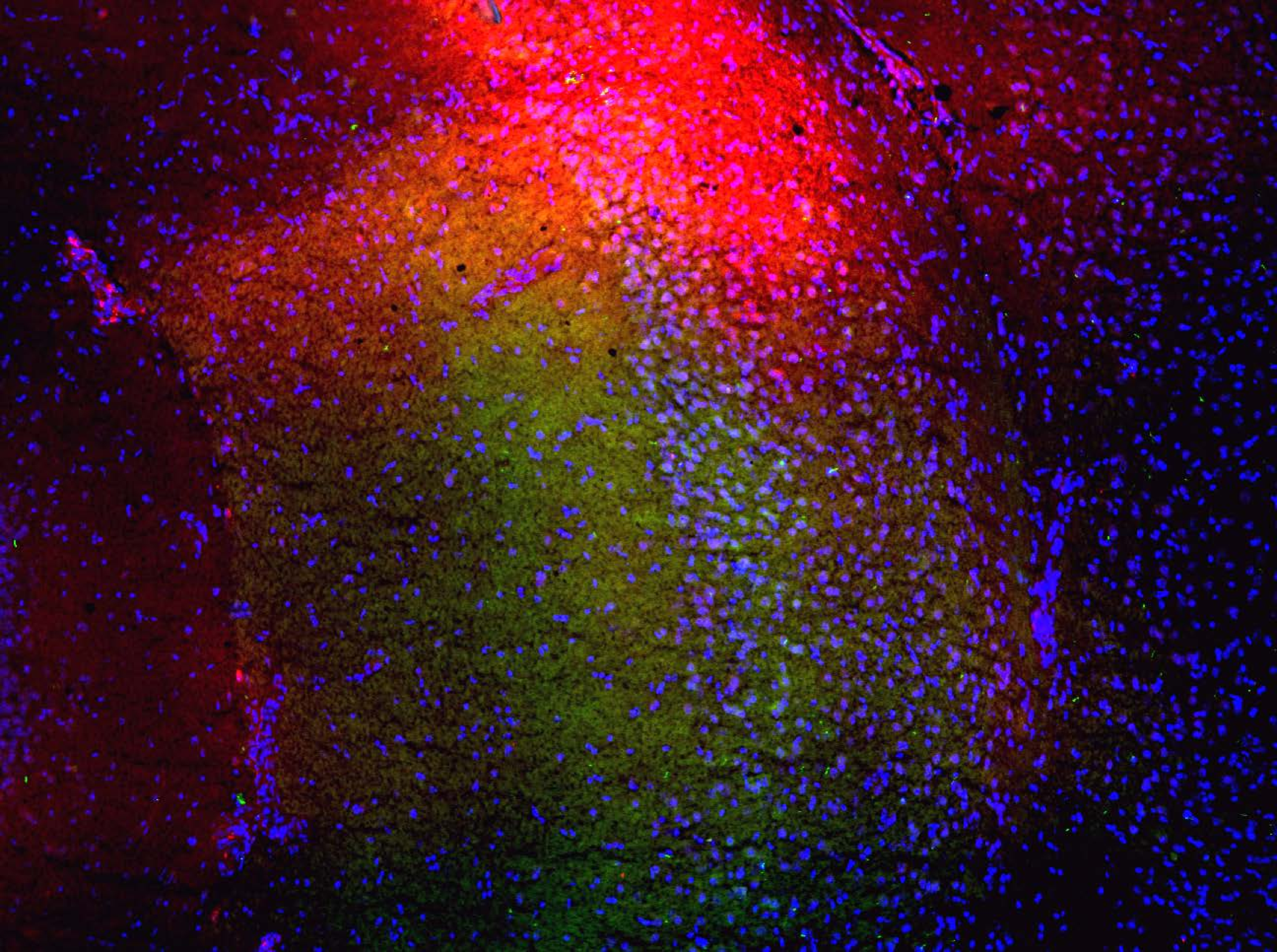

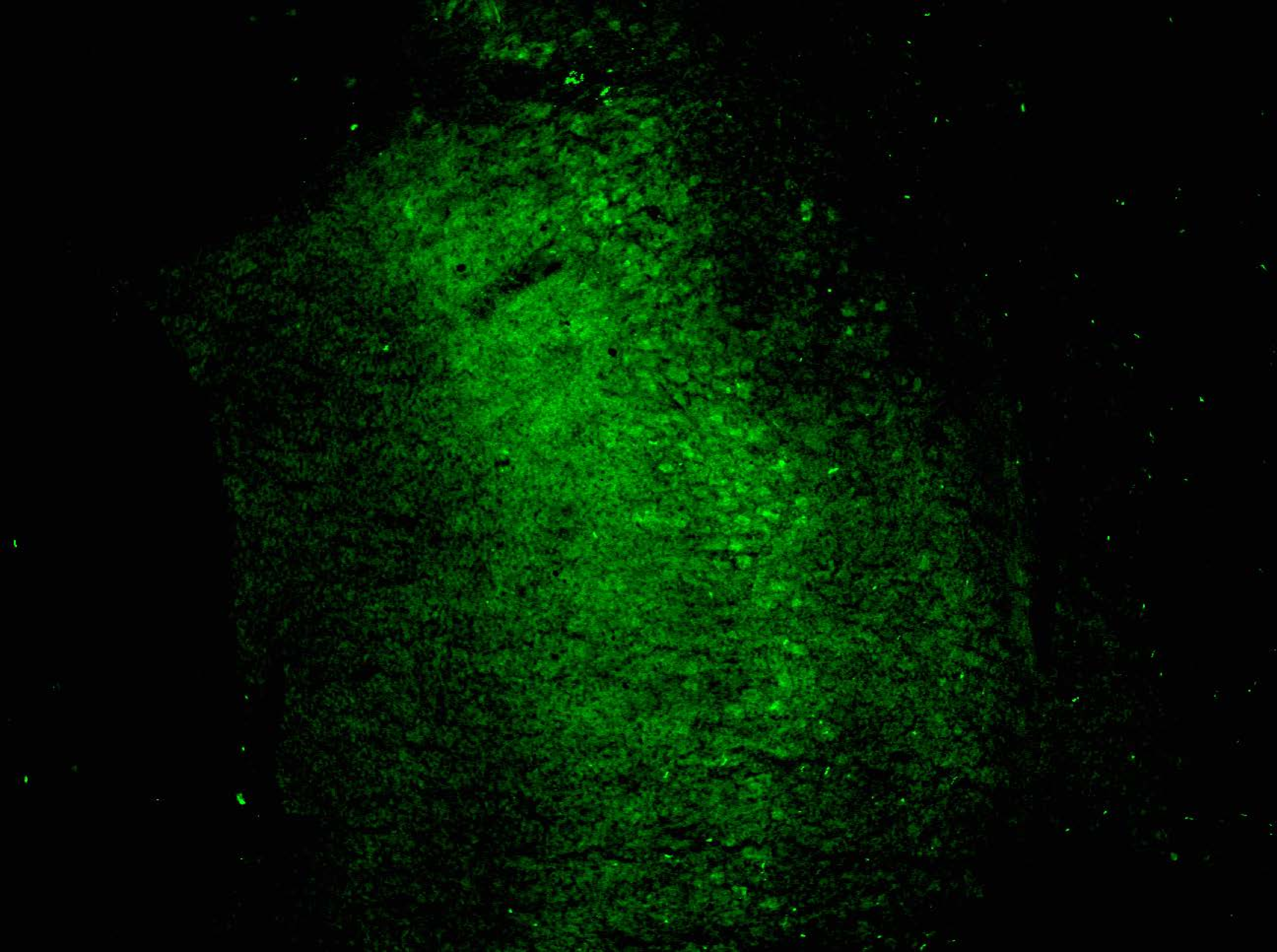

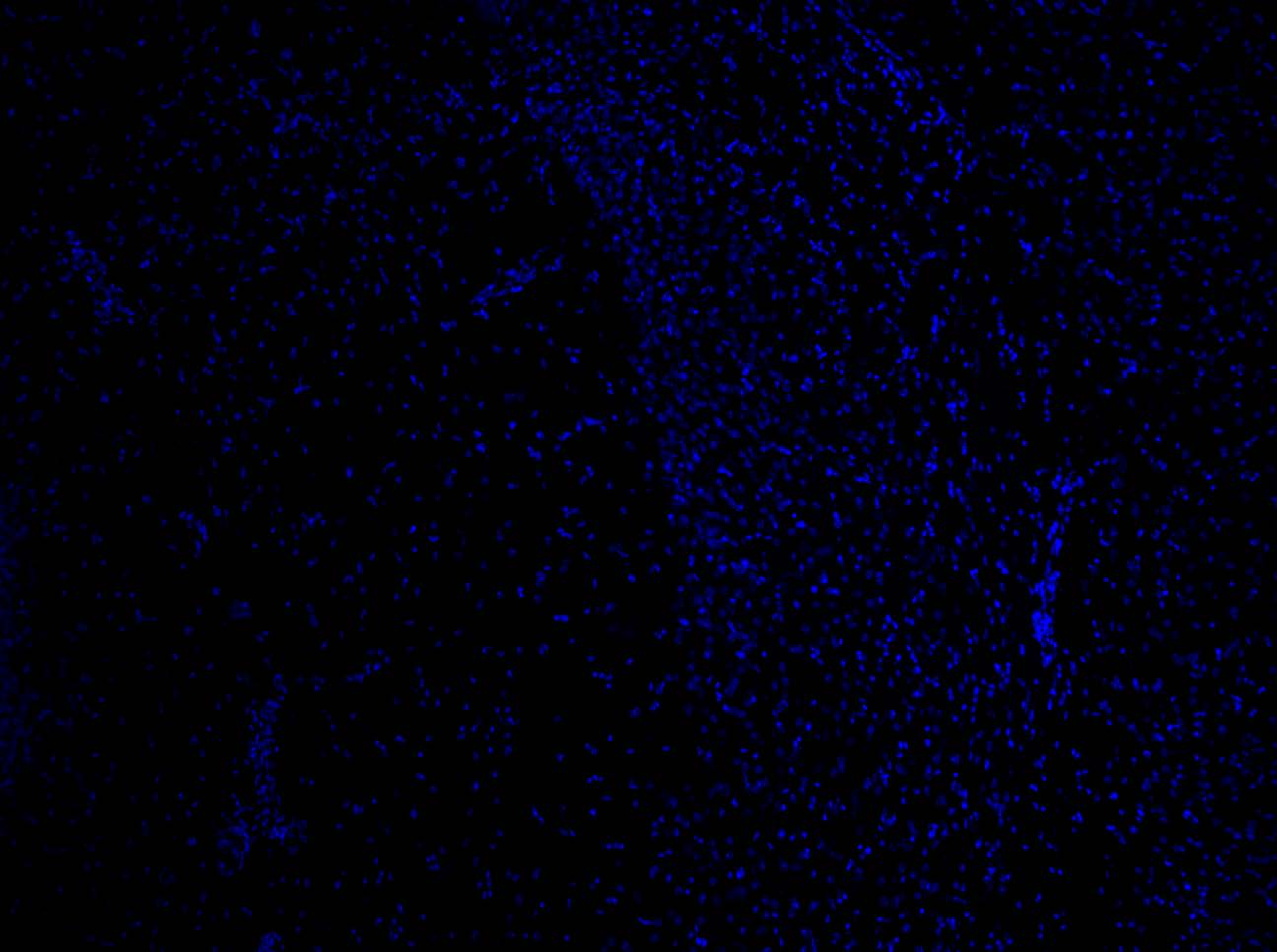

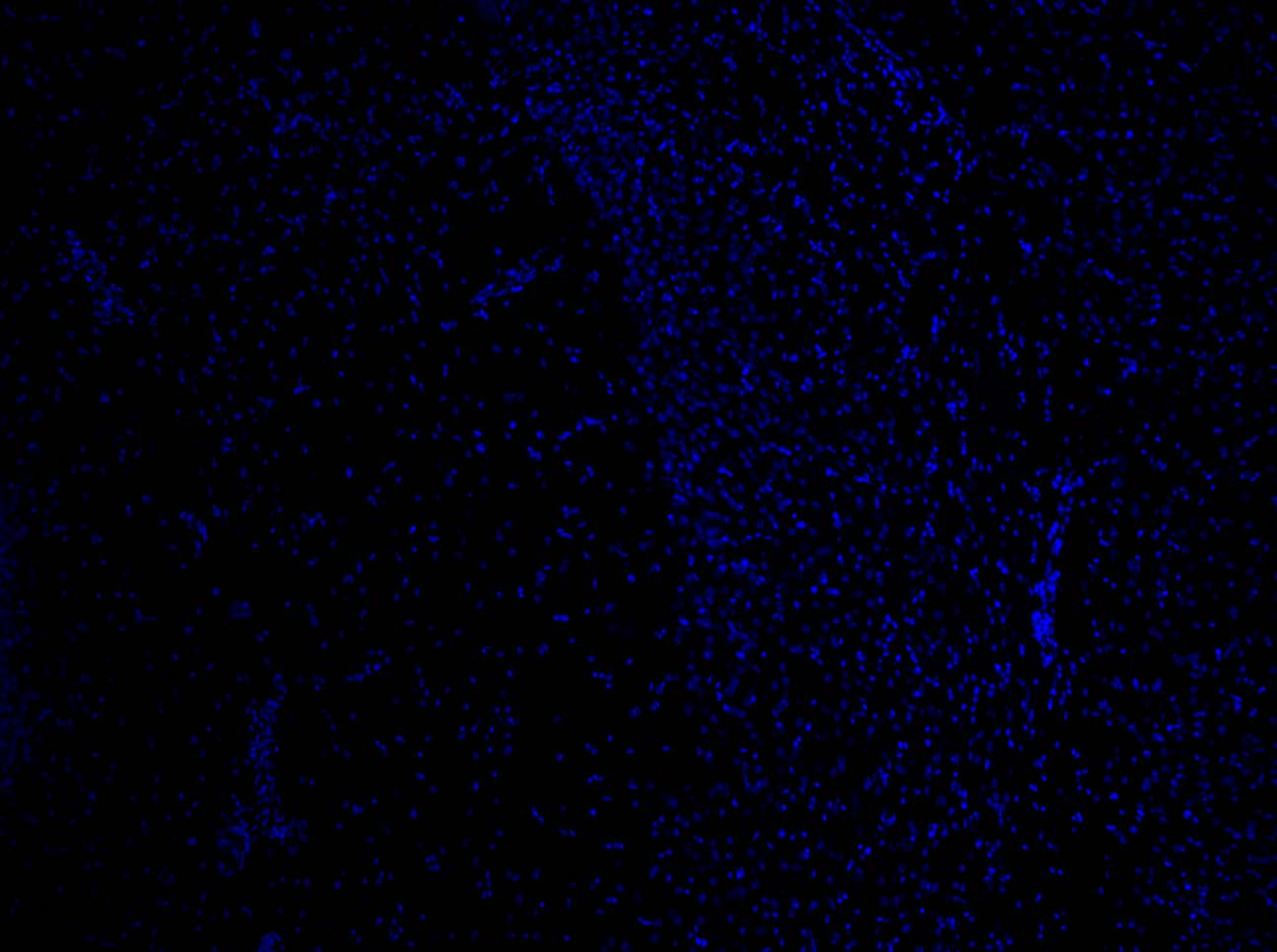

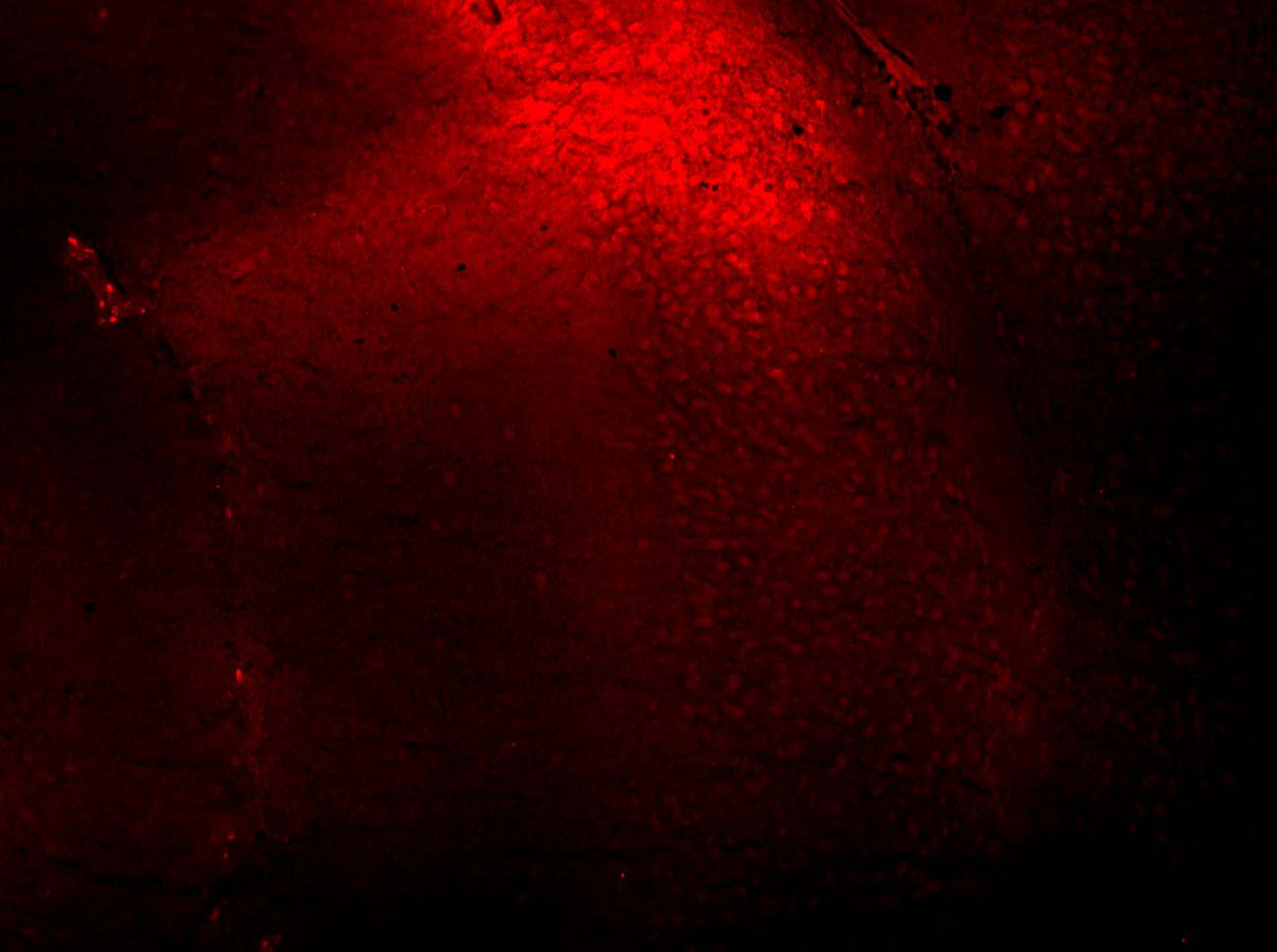

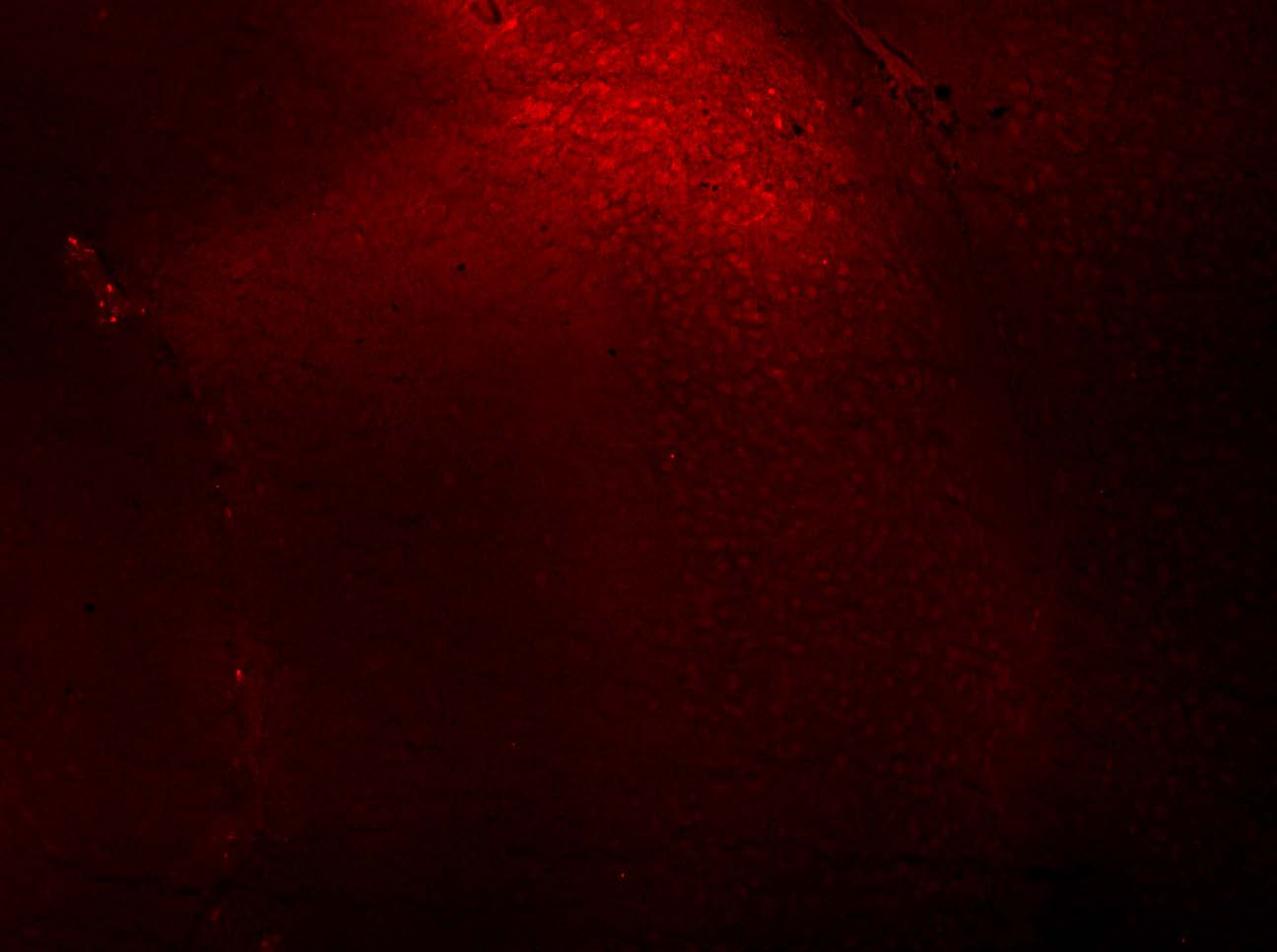

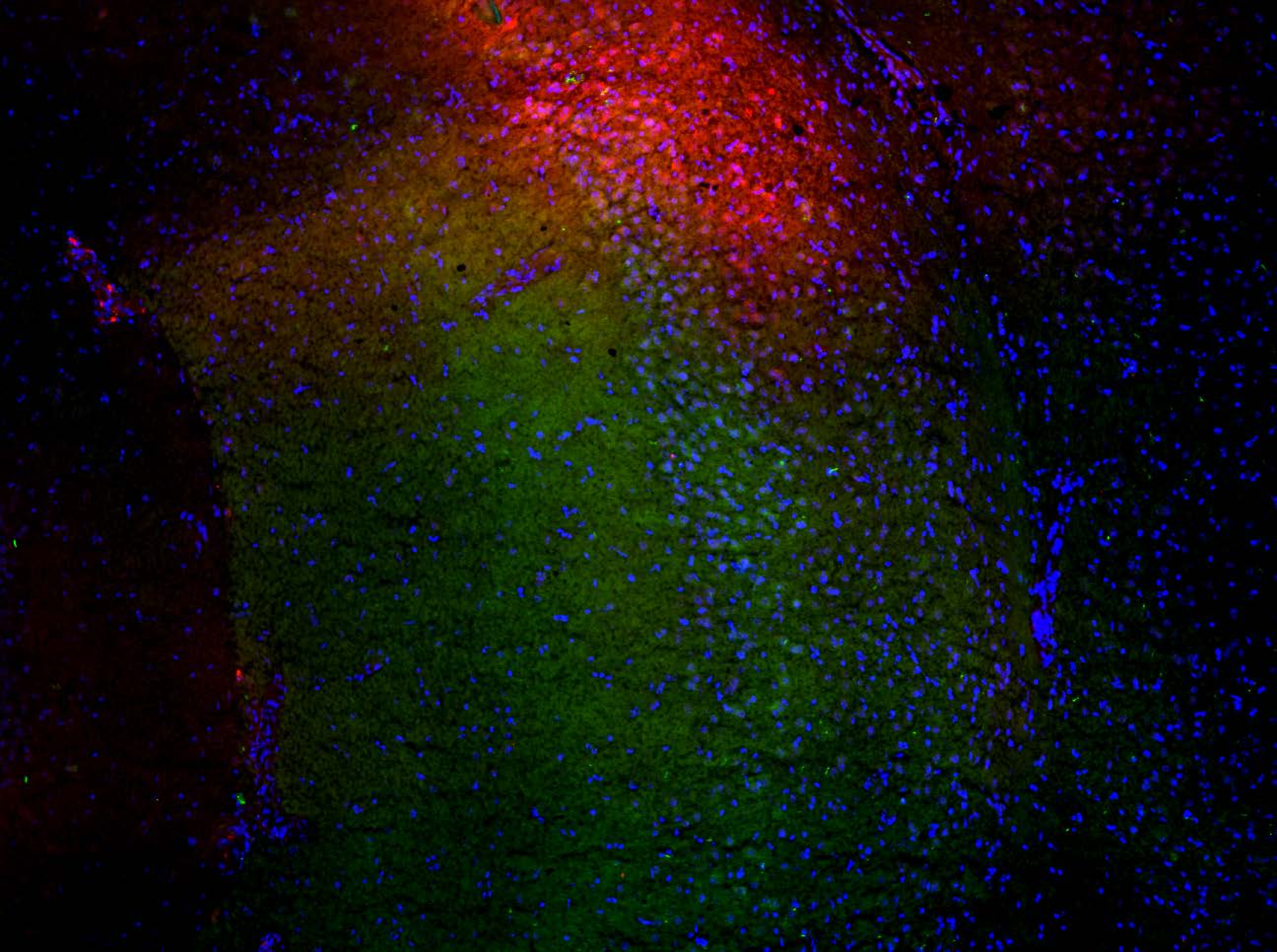

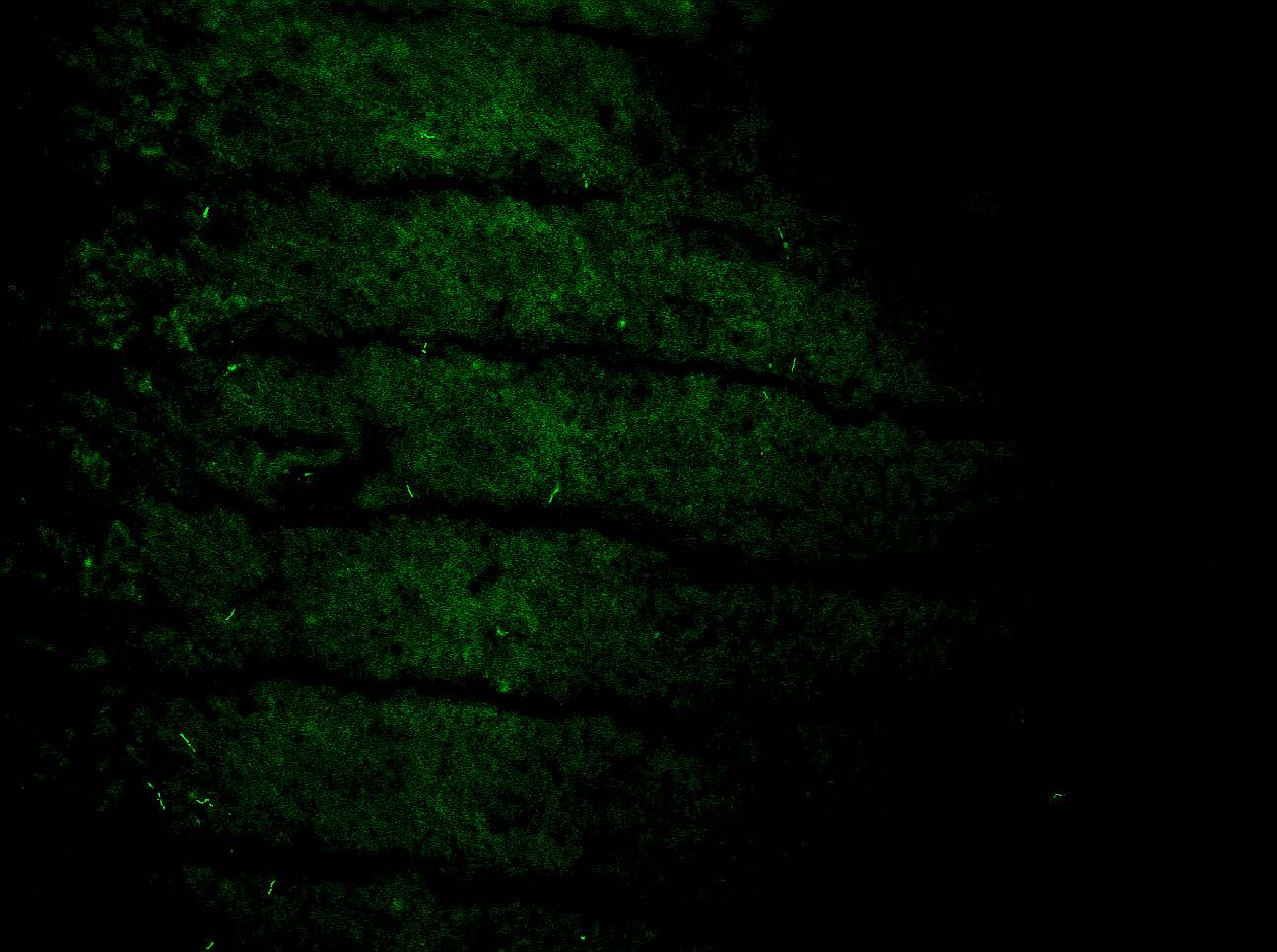

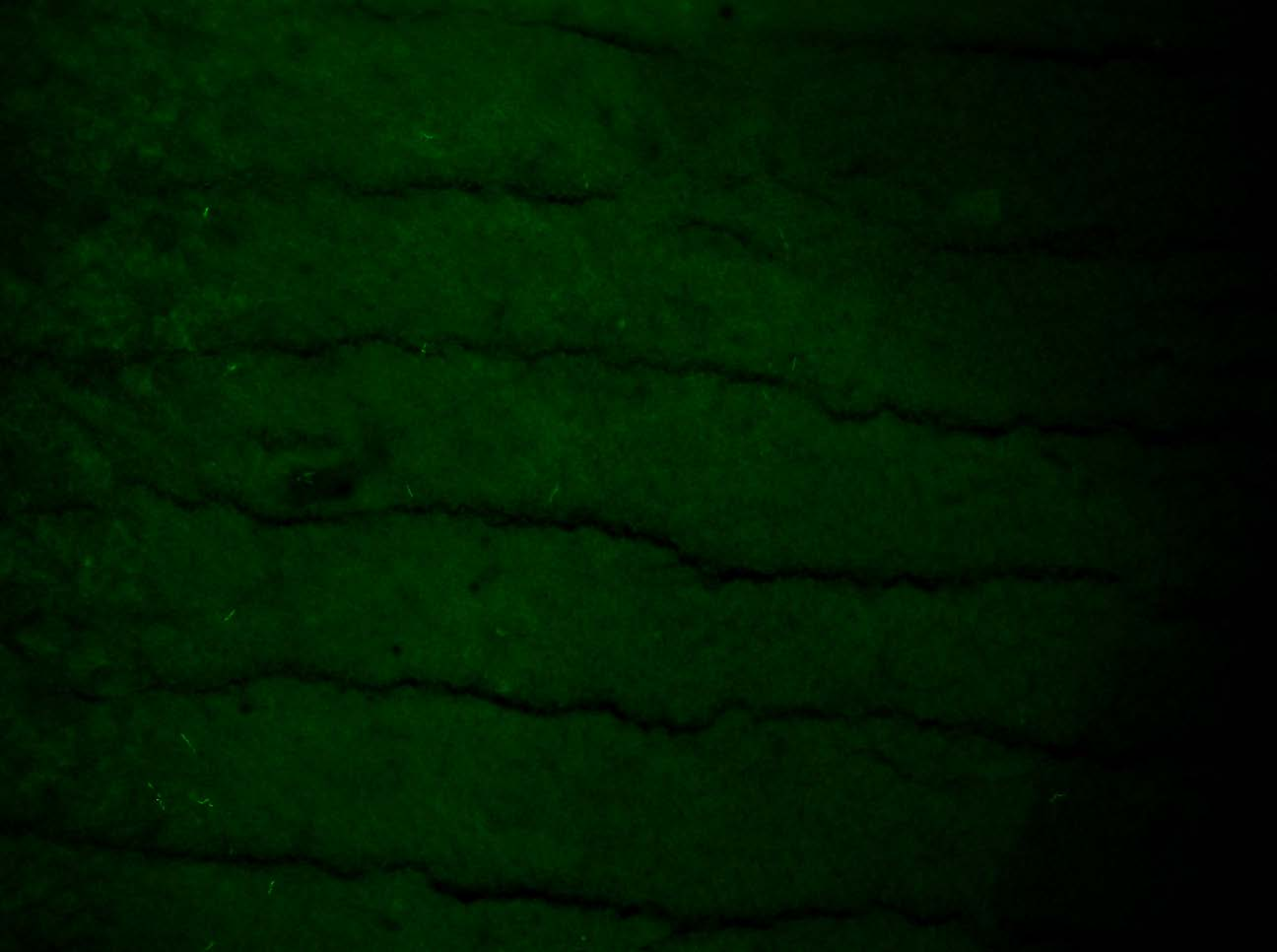

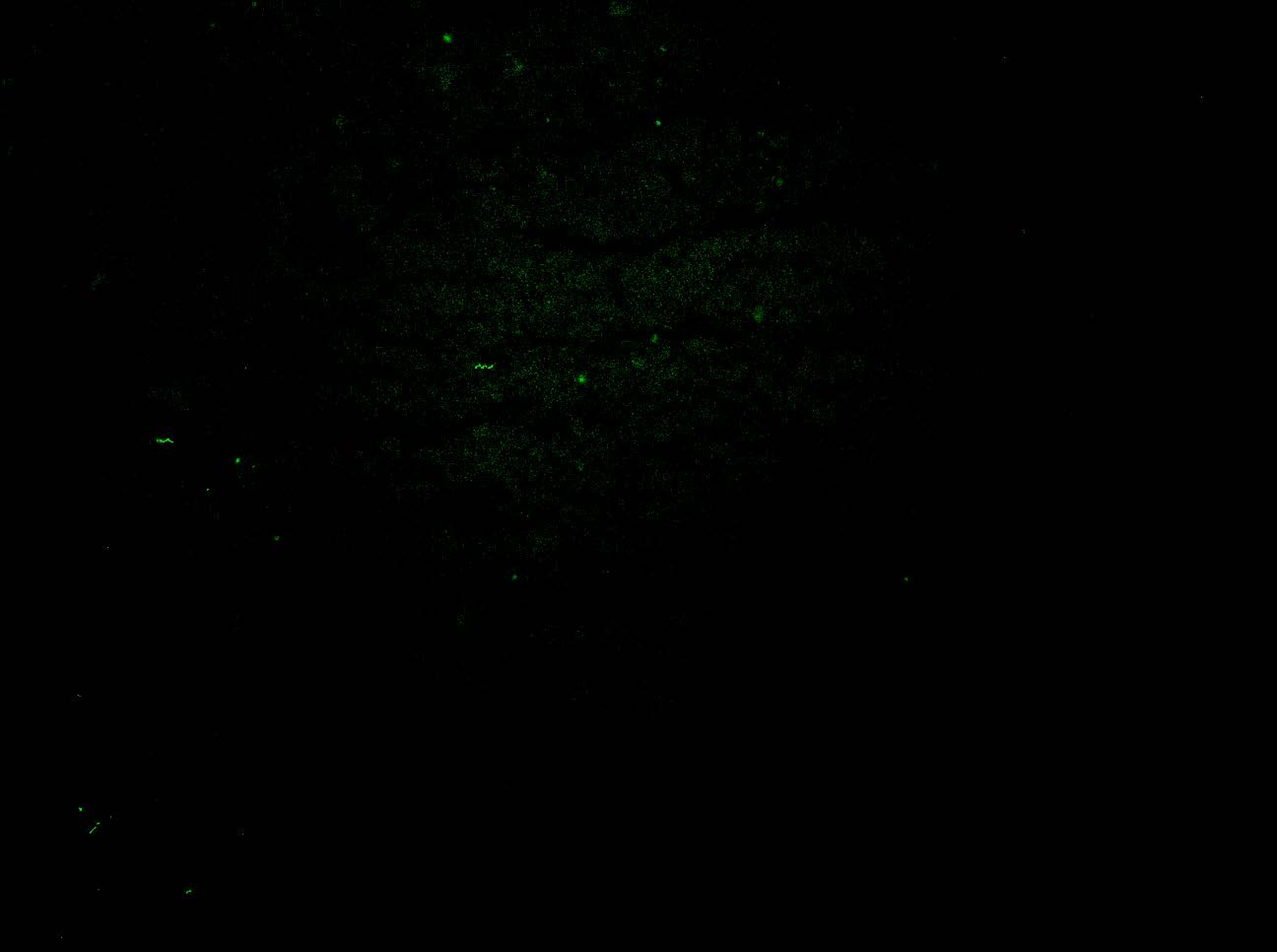

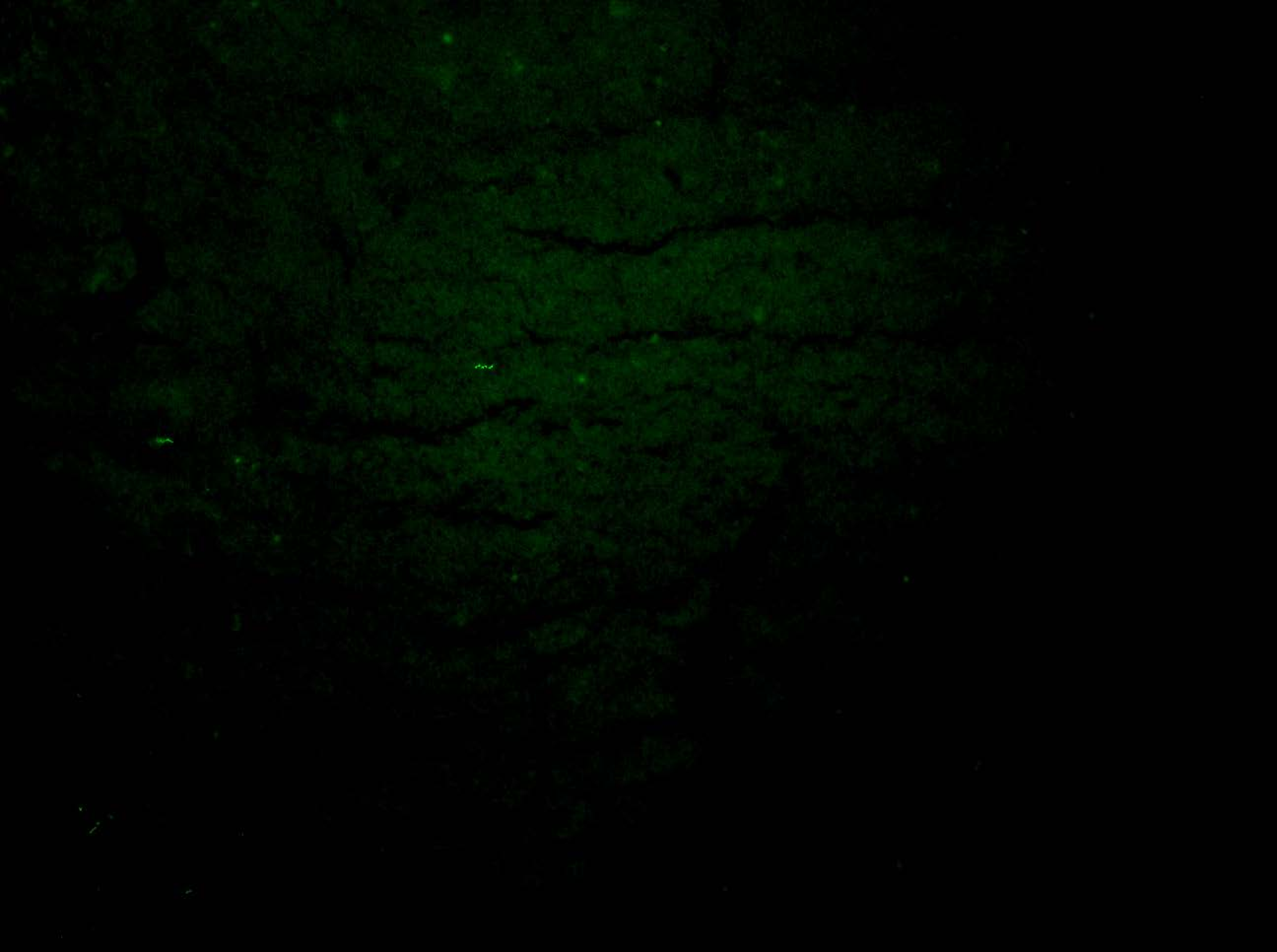

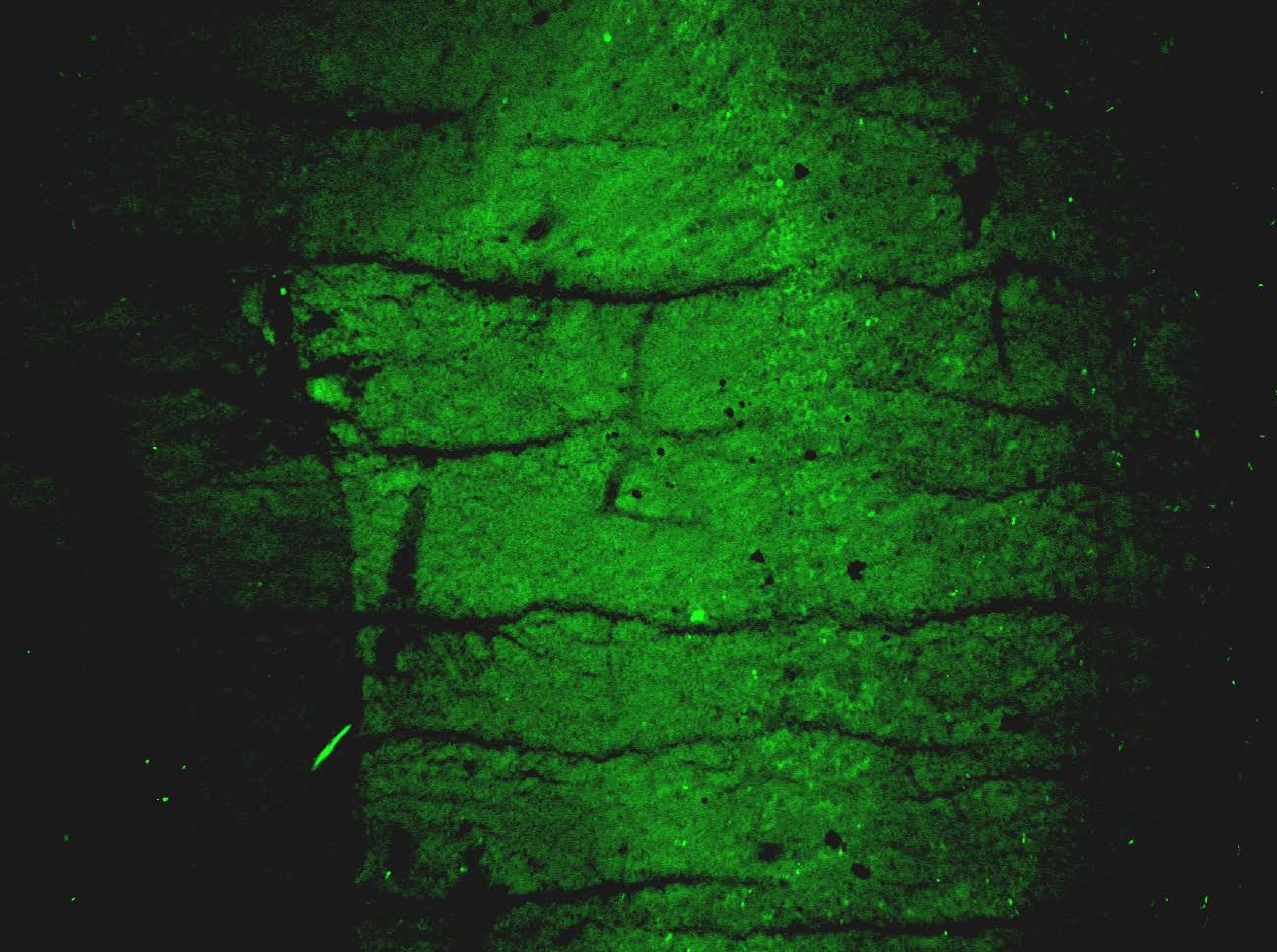

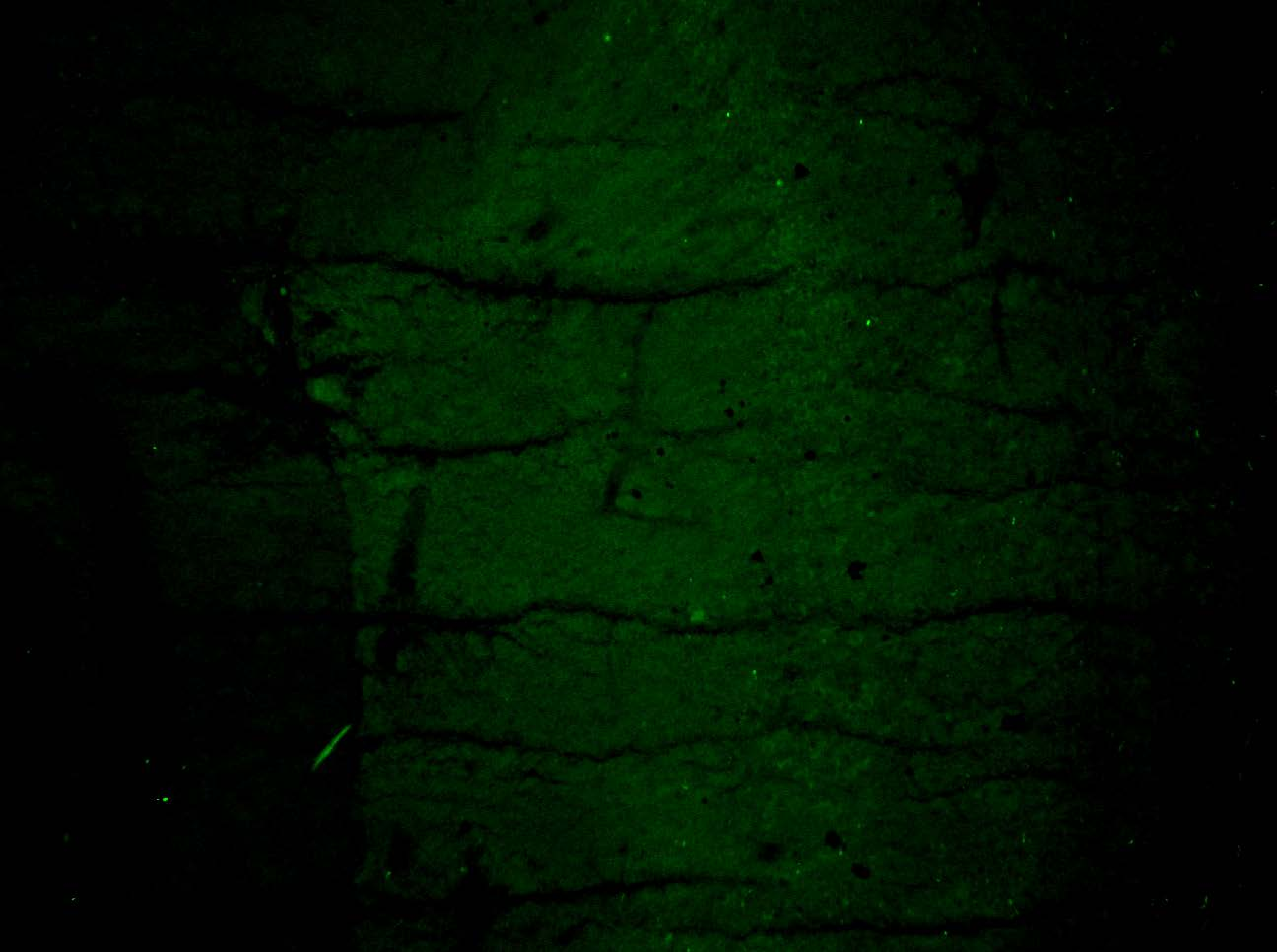

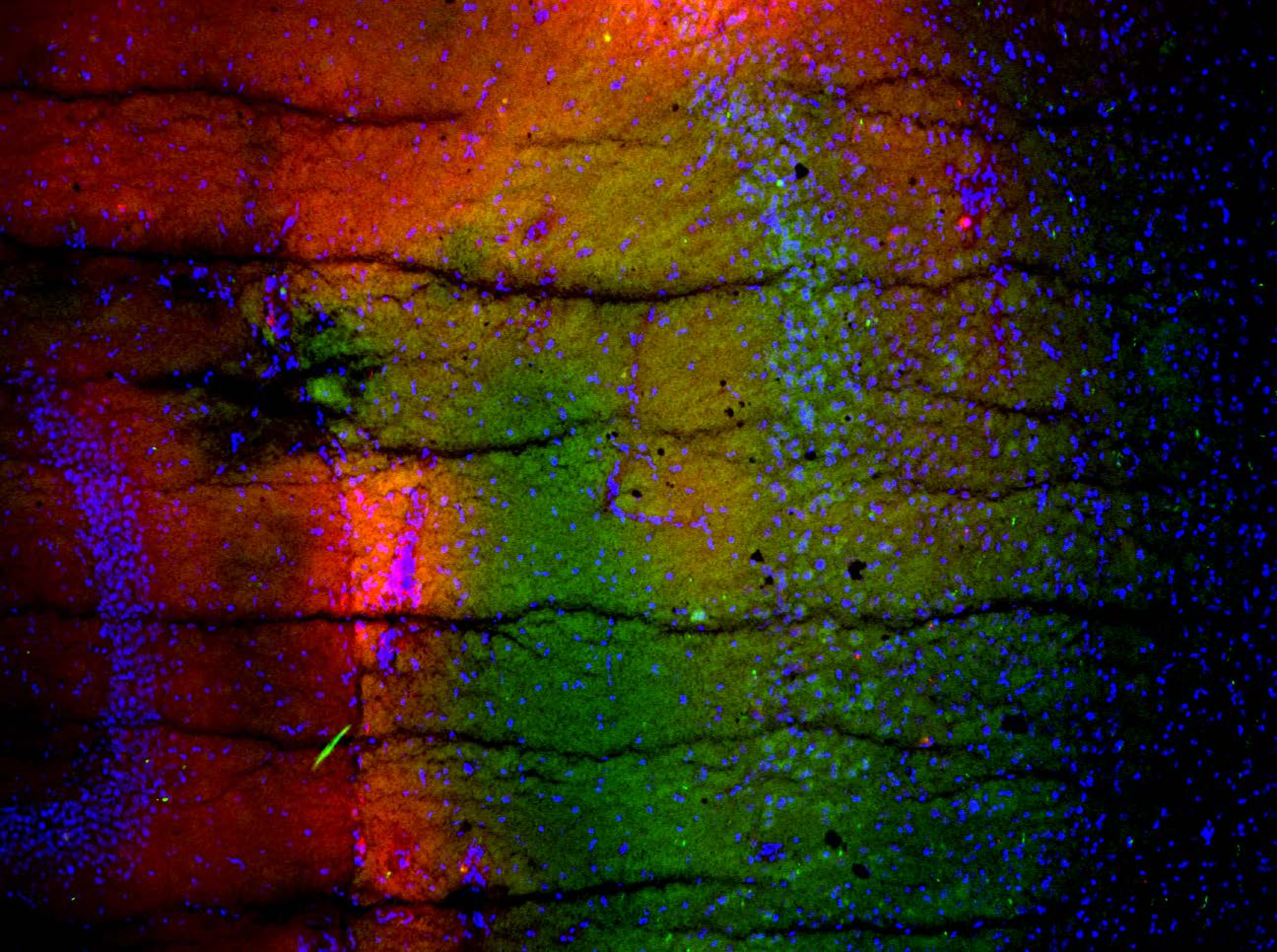

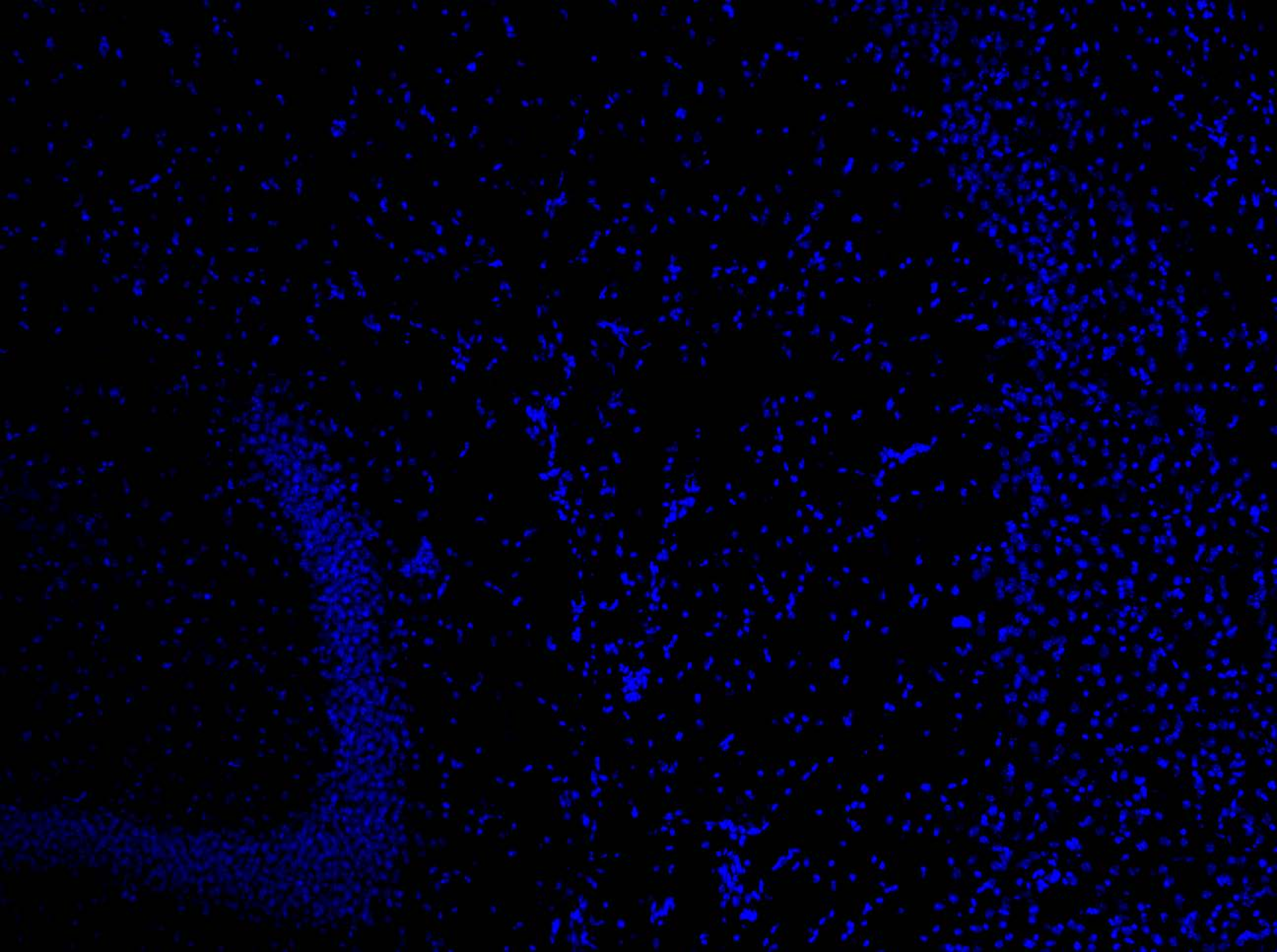
